# Structural assembly of the human Miro1/2 GTPases based on the crystal structure of the N-terminal GTPase domain

**DOI:** 10.1101/729251

**Authors:** Kyle P. Smith, Pamela J. Focia, Srinivas Chakravarthy, Eric C. Landahl, Julian L. Klosowiak, Sarah E. Rice, Douglas M. Freymann

**Affiliations:** Department of Cell & Molecular Biology, Feinberg School of Medicine, Northwestern University, 303 East Chicago Avenue, Chicago, IL, 60611, USA; Department of Biochemistry & Molecular Genetics, Feinberg School of Medicine, Northwestern University, 303 East Chicago Avenue, Chicago, IL, 60611, USA; Biophysics Collaborative Access Team, Advanced Photon Source, Argonne National Laboratory, Bldg. 435B/Sector 18, 9700 S. Cass Avenue, Argonne, IL 60439, USA; Department of Physics, DePaul University, Chicago, IL, 60614, USA; Department of Physical Therapy and Human Movement Sciences, Feinberg School of Medicine, Northwestern University, Chicago, IL, 60611, USA

**Keywords:** Miro, RhoT, Gem1p, GTP-binding protein, mitochondrial dynamics, crystal structure

## Abstract

Dysfunction in mitochondrial dynamics is believed to contribute to a host of neurological disorders and has recently been implicated in cancer metastasis. The outer mitochondrial membrane adapter protein Miro functions in the regulation of mitochondrial mobility and degradation, however, the structural basis for its roles in mitochondrial regulation remain unknown. Here, we report a 1.7Å crystal structure of N-terminal GTPase domain (nGTPase) of human Miro1 bound unexpectedly to GTP, thereby revealing a non-catalytic configuration of the putative GTPase active site. We identify two conserved surfaces of the nGTPase, the “SELFYY” and “ITIP” motifs, that are potentially positioned to mediate dimerization or interaction with binding partners. Additionally, we report small angle X-ray scattering (SAXS) data obtained from the intact soluble HsMiro1 and its paralog HsMiro2. Taken together, the data allow modeling of a crescent-shaped assembly of the full-length soluble domain of HsMiro1/2.

**PDB reference:** Crystal structure of the human Miro1 N-terminal GTPase bound to GTP, 6D71

## 1. Introduction

Mitochondrial motility (Schuler et al., 2017; Sheng and Cai, 2012; Caino et al., 2016) and other dynamics (Murley and Nunnari, 2016) are critically important in neuronal function, and their dysfunction is a hallmark in neurodegenerative diseases (Sheng and Cai, 2012; Li et al., 2004; Burté et al., 2015). Recent evidence suggests an emerging role in cell migration and possibly cancer metastasis as well (Murley and Nunnari, 2016; Schuler et al., 2017; Caino et al., 2016). Miro (also called RhoT), a type 2 transmembrane protein of the mitochondrial outer membrane, serves as an adapter of mitochondrial motility via the Kinesin-1/Dynein/TRAK machinery (Glater et al., 2006; MacAskill et al., 2009a). Knockout of Miro causes mis-localization and mitochondrial inheritance phenotypes on a cellular level (Chung et al., 2016) and knockout of Miro1 is embryonic lethal in mouse models (Nguyen et al., 2014; López-Doménech et al., 2016). In *D.melanogaster* Miro has been shown to be required for both antero and retrograde transport along microtubules (Guo et al., 2005; Russo et al., 2009). Miro is involved in actin-based movement through Myosin19 as well (Oeding et al., 2018; López-Doménech et al., 2018).

Miro comprises two GTPase domains (the nGTPase and cGTPase) which surround a pair of calcium-binding EF-hand domains. A C-terminal transmembrane peptide (TMD) mediates membrane attachment (Klosowiak et al., 2013) (**Fig. 1A**). Based on sequence signatures within the nGTPase domain and its apparent localization (Okumoto et al., 2017) the protein was initially termed a ‘rho’ GTPase (‘Mitochondrial Rho GTPase’, hence Miro), however, it has since become clear that the Miro proteins, which generally lack the ‘rho insert’ characteristic of the Rho GTPases, likely comprise a distinct subfamily of small GTPases (Wennerberg and Der, 2004; Koshiba et al., 2011). Miro is evolutionarily conserved down to *S.cerevisiae* (Frederick et al., 2004). In humans two paralogs of Miro (HsMiro1 and HsMiro2), which overall have ∼60% sequence identity, both play a role in mitochondrial dynamics (Fransson et al., 2006). In yeast, the single Miro homolog is termed Gem1p, with ∼30% overall sequence identity to the human Miro1 (Frederick et al., 2004).

**Figure 1.**
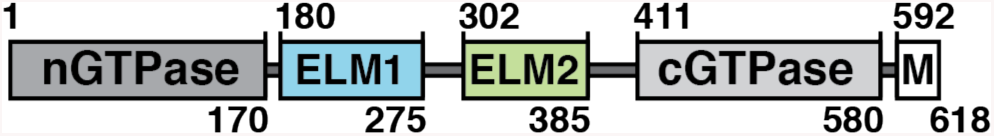
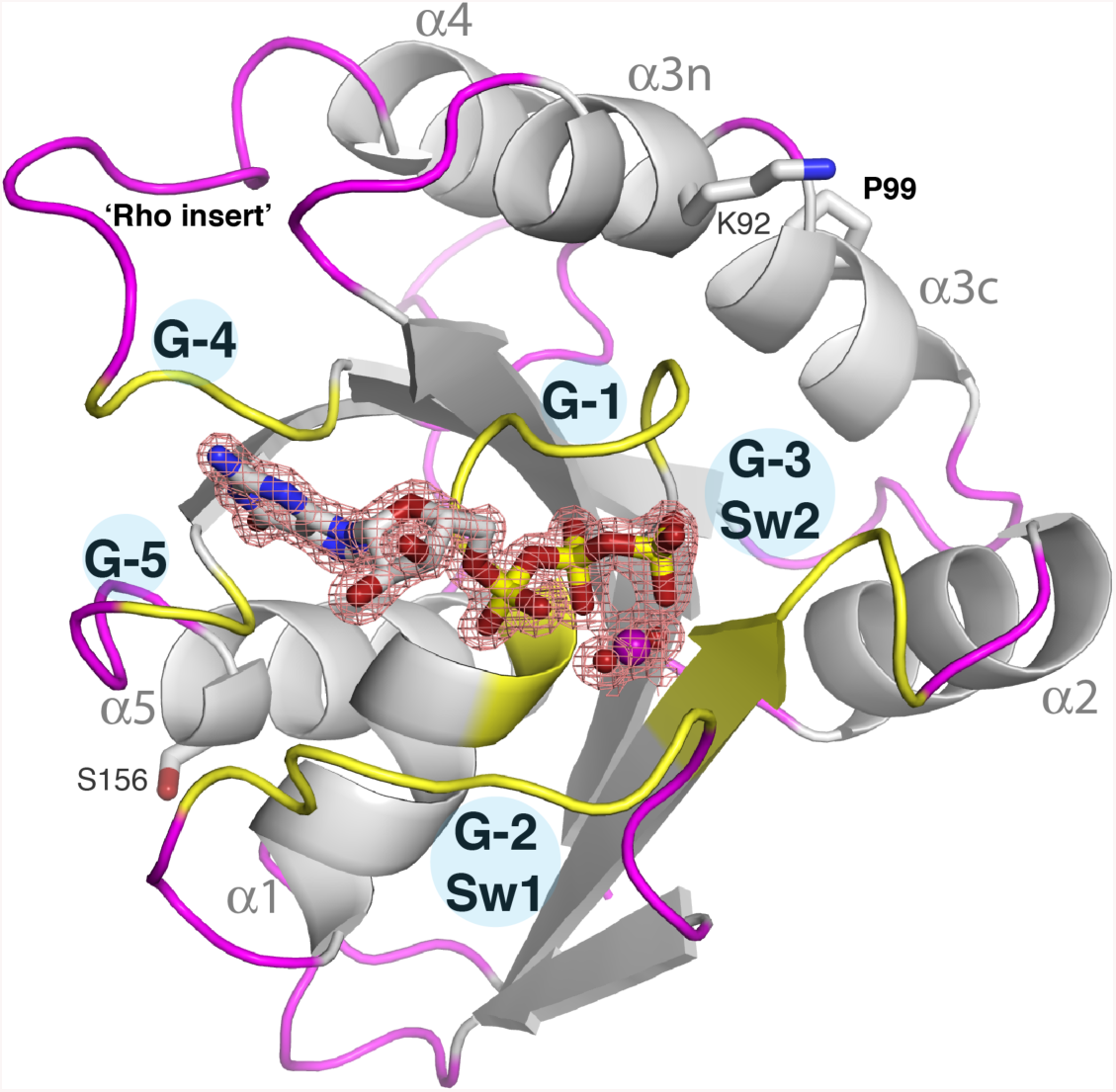
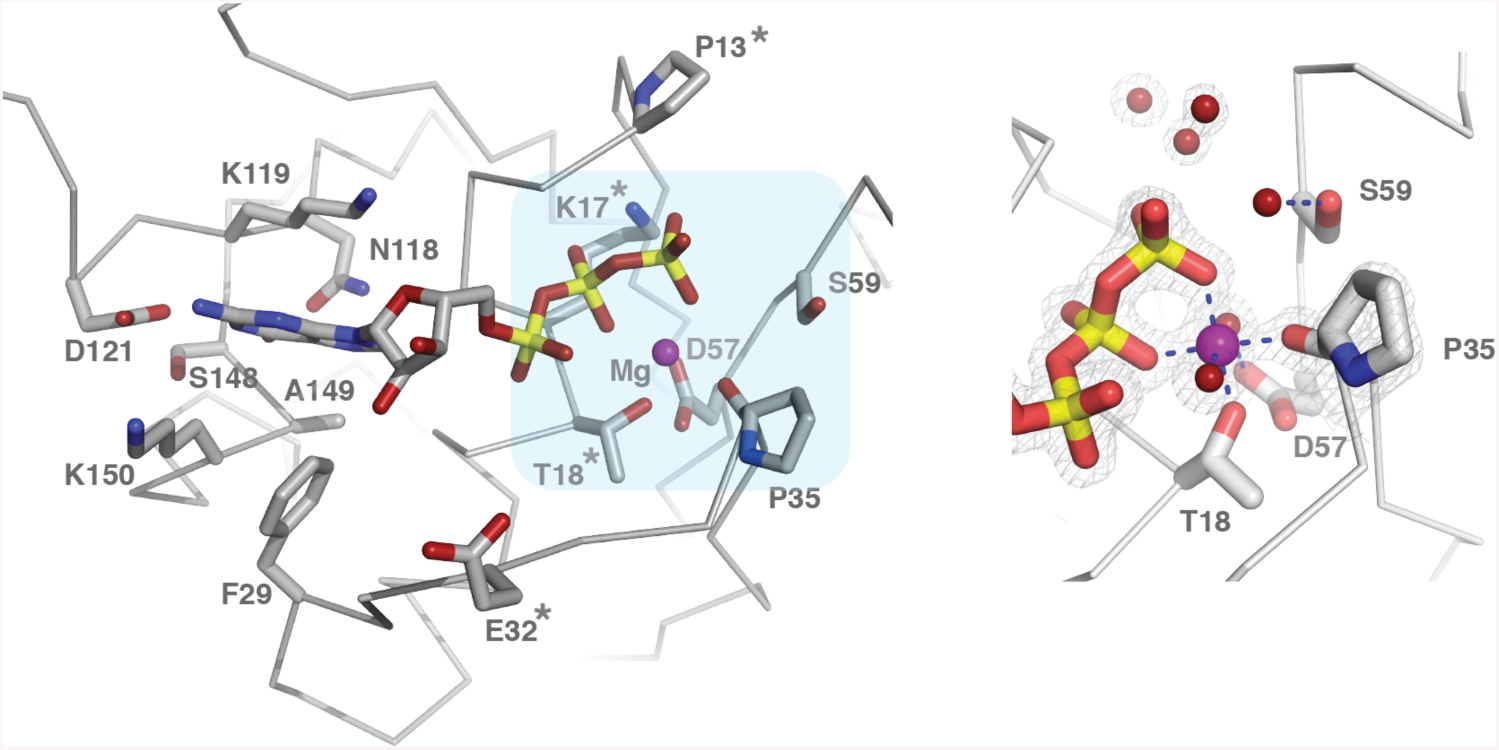
Crystal structure of the HsMiro1 nGTPase domain. (A) Bar diagram of the Miro domain architecture. Numbered residues indicate the limits of structured domains. ELM indicates the EF hand-Ligand mimic pairs. M indicates the transmembrane domain. (B) The overall fold of the HsMiro1 nGTPase domain. Guanine nucleotide binding regions (G-1 through G-5) are colored yellow and labeled. The Switch 1 (G-2) and Switch 2 (G-3) peptides are labeled as are helices α1-α5. The bound GTP is shown as sticks while the magnesium ion is shown as a purple sphere. The electron density of the bound Mg^2+^*GTP is shown contoured at 1.8σ. Pro99, which introduces a kink in helix α3 that is in common with other Rho GTPases, and the location of the ‘rho insert’ absent after G-4, are indicated. The sidechains of Ser156 and Lys92 indicate the positions of two regulatory post-translational modifications of the Miro nGTPase - PINK1 mediated phosphorylation at Ser156 (Wang et al., 2011; Shlevkov et al., 2016), and in mouse, acetylation at Lys107 (Kalinski et al., 2019). (C) **(left)** Overview of the active site interactions with bound GTP. Conserved residues that interact directly with GTP are labeled and shown as sticks. The positions of the ‘canonical’ GTPase residues that have been mutated in studies of Miro and Gem1p (Pro13, Lys17, Thr18, Glu32 (Thr33 in Gem1p)) are noted with ‘*’. The light blue area highlights the region shown in closeup. **(right)** A limited region of the 2Fo-Fc electron density is shown, contoured at 1.8 σ, showing the well-defined coordination interactions with the bound Mg^2+^ ion and the water structure at the active site. The carbonyl oxygen of Pro35 coordinates directly the active site Mg^2+^ ion. Water molecules at the putative active center, associated with Ser59, are shown as red spheres. Note that the water molecule interacting with Ser59 side chain hydroxyl is weakly bound (poorly defined in the map). Figures generated using PyMol (DeLano, 2002). A diagrammatic representation of the ligand interactions is shown in **Suppl. Fig. 2**.

The two EF-hand and C-terminal GTPase domains of Miro form a rigid assembly, termed ‘MiroS’ (Klosowiak et al., 2013; 2016). The EF-hand domains comprise canonical Ca^2+^ binding motifs paired with ‘hidden’ <u>E</u>F-hands that each interact with a putative ligand-<u>m</u>imic helix, hence are termed ELM1 and ELM2 (Klosowiak et al., 2013). While the cGTPase domain is structurally reminiscent of other small GTPases, structures of the human and drosophila cGTPase determined in the apo state or with GDP or GMPPCP bound reveal an open nucleotide binding site that exhibits no significant structural or functional change (Klosowiak et al., 2013; 2016). While mutagenesis studies clearly support a role for calcium binding to the ELM domains and nucleotide binding to the cGTPase domain in the regulation of Miro function (Fransson et al., 2006; MacAskill et al., 2009b; a; Saotome et al., 2008; Liu and Hajnóczky, 2009), the structural mechanisms by which these ligands modulate its activity remain unknown, and their specific roles remain controversial (Wang et al., 2011; Nguyen et al., 2014; Chang et al., 2011).

The nGTPase domain of Miro is functionally and structurally distinct from the cGTPase, and it is thought that the N-terminal GTPase domain of Miro may be more important for function, as mutations in the N-terminal domain appear to show more robust phenotypes than mutations in the C-terminal domain. For example, it was initially shown that only mutations of the nGTPase domain collapsed the mitochondrial network (Fransson et al., 2006). Mutation of corresponding residues of the putative GTPase active sites of the N- and C-GTPase domains likewise demonstrate dissimilar behaviors. For example mutation of T18N at the putative nGTPase active center is non-functional in recruiting CENP-F to mitochondria, while the corresponding S432N mutation of the cGTPase behaves like WT (Kanfer et al., 2015). And while both GTPase domains are required for function in yeast Gem1p, only mutations in the nGTPase, and not the cGTPase, alter mammalian mitochondrial transport into axons and dendrites (Guo et al., 2005). Furthermore, a recent study demonstrated acetylation of the equivalent residue to Lys92 in humans (mouse Lys105) in the nGTPase domain regulated axonal motility (Kalinski et al., 2019). On this basis it has been suggested that the nGTPase of Miro has the profound role in its function (Babic et al., 2015) while the cGTPase has a more ‘limited’ role (Koshiba et al., 2011). These observations may additionally explain why the nGTPase shows greater sequence conservation throughout evolution compared to the cGTPase.

Absent structural information about the nGTPase, dissection of the function of Miro *in vivo* has generally exploited mutations that are based on inferred functional relationships with the Ras family of GTPases (Fransson et al., 2003; Frederick et al., 2004; Saotome et al., 2008; Wang and Schwarz, 2009; MacAskill et al., 2009a; Koshiba et al., 2011; Kornmann et al., 2011; Kanfer et al., 2015; Babic et al., 2015). Such mutations have largely been interpreted as generating what have been termed ‘GTP-bound’ and ‘GDP-bound’ (or ‘active’ and ‘inactive’) states (Saotome et al., 2008; Babic et al., 2015). However, as little is known about the actual structure/function relationships of the Miro proteins, whether or not such mutations selected on the basis of analogy impact Miro GTPase function as anticipated remains essentially unknown. Indeed, it is becoming clear that the GTPase fold can be exploited in different ways by different biological systems, with the canonical ‘GTPase switch’ only one of several mechanisms by which GTPase domains can function (e.g. serving as a module for nucleotide-dependent assembly (Chappie et al., 2010; Focia et al., 2004; Yan et al., 2018)), and, therefore, that the biological response of a particular mutation can not necessarily be predicted. Additionally, the limited scope of the mutational library has restricted focus towards function of the Miro GTPase domain as a canonical GTPase switch, whereas clearly a complex set of additional interactions must be required in order for Miro to serve as a tether for regulation of mitochondrial dynamics.

Although the existence of a nucleotide hydrolysis activity by Miro has been inferred from its relationship to other members of the GTPase superfamily, it has received limited biochemical characterization, with the most extensive work carried out using the yeast homolog Gem1p. Indeed, the purified Gem1p readily hydrolyzes GTP, and mutational studies locate the activities to both its N- and C-GTPase domains (Koshiba et al., 2011). The purified *Drosophila* DmMiro has been shown to hydrolyze GTP, as well (Lee et al., 2016). That observations based on Gem1p can be readily extended to the mammalian Miro remains controversial (Kornmann et al., 2009; Lewis et al., 2016; Nguyen et al., 2012). However, recently, biochemical characterization of HsMiro1/2 domain constructs corresponding to the nGTPase, cGTPase, and MiroS has confirmed that both putative GTPase domains exhibit readily measurable GTP nucleotide hydrolysis activity *in vitro* (Peters et al., 2018). The rate constants for hydrolysis for both domains towards GTP are similar to that of the Ras GTPase. Remarkably, however, the activity of the cGTPase was found to be promiscuous, hydrolyzing ATP and CTP in addition to GTP. The authors propose a structural basis for these differences based on modeling studies, which have yet to be confirmed (Peters et al., 2018). However, it seems clear that regulation of nucleotide binding and hydrolysis by both domains likely play a key role in Miro function.

In order to fully understand the regulatory mechanisms and functional interactions of Miro it will be essential to structurally characterize each of its domains in the context of the full-length protein and to identify the contacts that mediate its intra- and intermolecular assemblies. To that end, we have carried out the X-ray crystallographic and solution scattering studies of the human Miro1 and Miro2 described here. We seek to understand the functional role of the remarkable domain organization of Miro and to dissect the mechanisms by which its interactions contribute to regulation of mitochondrial dynamics. Previously, we determined the crystal structures of the DmMiro and HsMiro1 EF-hands and cGTPase domains (Klosowiak et al., 2013), and the structure of the HsMiro2 cGTPase domain (Klosowiak et al., 2016). Here, we have determined the crystal structure of the HsMiro1 nGTPase domain bound, unexpectedly, to un-hydrolyzed GTP. The structure allows us to locate two highly conserved surfaces of the nGTPase domain distinct from the active site which mediate packing contacts in the crystal and that may be poised for functional interaction. In conjunction with small angle X-ray scattering (SAXS) data obtained from both HsMiro1 and HsMiro2, we propose a model for the structure of the entire extra-membranous domain (residues 1-592) of Miro.

## 2. Experimental

### 2.1. Cloning and Molecular Biology

HsMiro2 cDNA were graciously provided by the Shaw lab (Fransson et al., 2006). Fragments were PCR amplified (CloneAmp HiFi PCR Premix, Clonetech) and subcloned into pET-28b vectors (Novagen) as 6xHis tagged proteins via Nco1 and Xho1 restriction sites using ligation-independent cloning (In-Fusion, Clonetech). Site-directed mutagenesis (Quikchange, Agilent) was employed to create all protein mutants using primers designed in-house (Integrated DNA Technologies). Residue number for the constructs are as follows: HsMiro1 full-length (1-592) c-terminal 6xhis tag (expected MW ∼68kD), HsMiro2 full-length (1-588) c-terminal 6xhis tag (expected MW ∼66kD), HsMiro1/2 nGTPase (1-180) n-terminal 6xhis tag. Full HsMiro1 (Q8IXI2), HsMiro2 (Q8IXI1), DmMiro (Q8IMX7), and Gem1p (P39722) amino acid sequences are accessible via Uniprot.

### 2.2. Protein Expression and Purification

#### HsMiro1 nGTPase

All recombinant proteins were expressed using *E. coli* BL21 RP (DE3) strain (Novagen). After inoculating 2 L of TPM media with 2 mL of overnight starter culture, the cells were grown with shaking at 200 rpm for approximately 5 hours at 37°C to OD_600_=0.45. The temperature was then lowered to 16°C for 1 hour and then cells were induced at OD_600_=0.90 with 0.125 mM IPTG for approximately 18 hours overnight. The cells were resuspended in 50 mM HEPES*HCl (pH 8.0), 500 mM NaCl, 1 mM MgCl_2_, 0.5 mM TCEP, 5% sucrose (w/v), and 0.05% Tween20 (v/v) before flash freezing in liquid nitrogen. Cell pastes were stored at −80°C before lysis and purification. Cells were thawed at 37 °C and all lysis and purification steps were performed at 4°C. The cell pastes were sonicated for 4×45 seconds before clarifying at 35,000 rpm for 45 min. The soluble fraction from 8 L of culture was incubated with 8 mL of 50% Co^2+^ TALON bead slurry, washed (8mM imidazole), then eluted with IMAC Elution Buffer (500 mM imidazole, 200mM NaCl), then slowly diluted 4-fold in Buffer A (25 mM HEPES*HCl pH 7.4, 1 mM MgCl_2_, 0.5 mM TCEP) to be loaded onto a 5mL HiTrap Q ion exchange column. The protein was eluted using a 18%-30% 1M NaCl step profile (Buffer A supplemented with 1M NaCl) and concentrated to approximately 2 mL using 10 kDa MWCO spin concentrators (Amicon) before running over a 16/60 Superdex 200 size exclusion column (GE Healthcare) in purification buffer (25 mM HEPES*HCl pH 7.4, 150 mM NaCl, 0.5mM MgCl_2_, 0.5 mM TCEP, 5% w/v sucrose). Peak fractions were pooled and stored at −80°C. Approximate final yields were ∼1 mg pure protein per 2 L of culture. Protein purity was assayed using SDS-PAGE.

#### HsMiro1 and HsMiro2 full-length

Expression, lysis and purification of the His-tagged proteins were carried out as described above, with the exception that 1mM CaCl_2_ was added to all buffers. Subsequently, samples were diluted into Buffer B (25mM Tris pH 8.5, 1mM MgCl2, 1mM CaCl2, 0.5mM TCEP, 2.5% sucrose) to be loaded onto a 5ml HiTrap Q column, followed by elution using a 8-50% gradient (Buffer B supplemented with 1M NaCl). Peak fractions were pooled and injected over a 16/60 S200 gel filtration column equilibrated with 25mM Tris 8.5, 500mM NaCl, 1mM MgCl2, 1mM CaCl2, 0.5mM TCEP, 2.5% sucrose. Peak fractions were concentrated to ∼2.5 mg/ml and flash frozen.

### 2.3. Crystallization and X-ray Data Collection

Protein was concentrated to ∼10 mg/mL in purification buffer (25 mM HEPES*HCl (pH 7.4), 150mM NaCl, 0.5 mM MgCl_2_, 0.5 mM TCEP, and 5% sucrose (w/v)). Crystallization trials were set up at 22 °C with the commercial sparse matrix screens JCSG+, PACT, PEGs, and PEGs II Suites (Qiagen) in INTELLI-WELL 96-3 LVR sitting drop vapor diffusion plates using a Crystal Phoenix robot (Art Robbins Instruments). 2:3, 1:1, and 3:2 protein:precipitant ratios were tested in a 1.0 µL total drop volume with a 90 µL reservoir. Rod-shaped crystals initially grew after 12 hours only in condition #38 from the PACT Suite (0.1 M MMT buffer pH 5, 25% (w/v) PEG 1500). Crystal diffraction quality was optimized using INTELLI-WELL 24-4 sitting drop vapor diffusion plates (Art Robbins Instruments) over a range of MMT Buffer pH 4.5-5.5 + PEG 1500 10-30% (w/v) concentrations, and testing different guanine nucleotide conditions. Crystals grew to about 30 µm with rod and cube-shaped morphologies. Cryo-protection was not required during data collection. Measurement of X-ray diffraction data was performed at 100K at the beamlines of Sector 21 of the Life Sciences Collaborative Access Team (LS-CAT) of the Advanced Photon Source (APS) in the Argonne National Laboratory (ANL). Data were measured on a MarMosaic 225 CCD detector and processed using Mosflm (Battye et al., 2011). 1 sec exposures were recorded every 1° of rotation over 120 ° at 65% beam attenuation.

### 2.4. Structure Determination and Refinement

An initial 3.2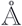 resolution dataset was obtained from a crystal grown in the presence of a mixture of 1mM GDP and (as subsequently discovered) GTP. The structure was determined by molecular replacement using the Phenix add-on Ensembler (Wang and Snoeyink, 2008) to create an overlapped model of GTPases based on HsMiro1 cGTPase (PDB ID=5KSP), RRas2 (PDB ID=2ERY), and RhoB (PDB ID=2FV8). The model obtained from Phenix Autobuild (Terwilliger et al., 2008) accounted for 82% of the polypeptide chain (304 of 372 residues in the asymmetric unit). Electron density corresponding to a bound nucleotide was clearly visible and could be unambiguously interpreted as arising from bound guanosine tri-phosphate, not the anticipated guanosine di-phosphate. Subsequently a 1.7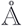 dataset was obtained from a crystal grown using protein at 10mg/ml supplemented with 1µM GDP, with a 1:1 2.0 µL drop volume over a reservoir of 0.6M MMT pH 5, 22% PEG 1500. This dataset also revealed the protein to be only GTP-bound (as did additional lower-resolution datasets from crystals obtained in the presence of 1mM, rigorously purified, GDP) and it was used for rebuilding and refinement of the final structure with COOT (Emsley et al., 2010) and phenix.refine (Afonine et al., 2012). The space group is P2_1_2_1_2_1_, with two molecules related by a non-crystallographic two-fold symmetry (NCS) axis in the asymmetric unit. Residues 4-175 (chain A) and 4-172 (chain B), as well as the bound Mg^2+^GTP ligands, were readily built into the electron density map; residues 1-3 and 176/173-180 were not visible and are presumed to be unstructured. Superposition of the NCS-related monomers yields an RMSD of 0.232 Å over 136 C*α* atoms. GTP ligand geometry restraints were added with Phenix eLBOW (Moriarty et al., 2009). Once the crystallographic statistics reached R_work_= 0.252 and R_free_ = 0.296, water molecules were added in every subsequent round of refinement. Final statistics are reported in **Suppl. Table 1**. PyMOL was used to generate all figures (DeLano, 2002). Superpositions and RMSD values were calculated using the SUPER routine in PyMOL.

### 2.5. Homology Modeling

Homology models of the Gem1p and DmMiro nGTPase domains were generated using I-TASSER (Zhang, 2009; Yang and Zhang, 2015), using the HsMiro1 nGTPase structure to provide the initial template. For Gem1p nGTPase the Iden1 (percentage sequence identity in the threading aligned region in the query sequence) with respect to HsMiro1 is 0.30, the C-score (the confidence score for estimating the quality of predicted models by I-TASSER) is 0.41 and the TM-score (a similarity score insensitive to the local modeling error (Zhang and Skolnick, 2004)) is 0.77. For DmMiro nGTPase Iden1 is 0.66, the C-score is −0.48, and the TM-score is 0.65. The residue-level quality of the protein structure predictions were calculated using ResQ (Yang et al., 2016) based on local variations of modeling simulations and the uncertainty of homologous alignments. These values, the ‘estimated local accuracy’, are plotted using the PyMol ‘Putty’ representation in **Fig. 2B**. Additional sequence alignments were carried out with T-COFFEE and EMBOSS-NEEDLE (Di Tommaso et al., 2010; Rice et al., 2000).

**Figure 2.**
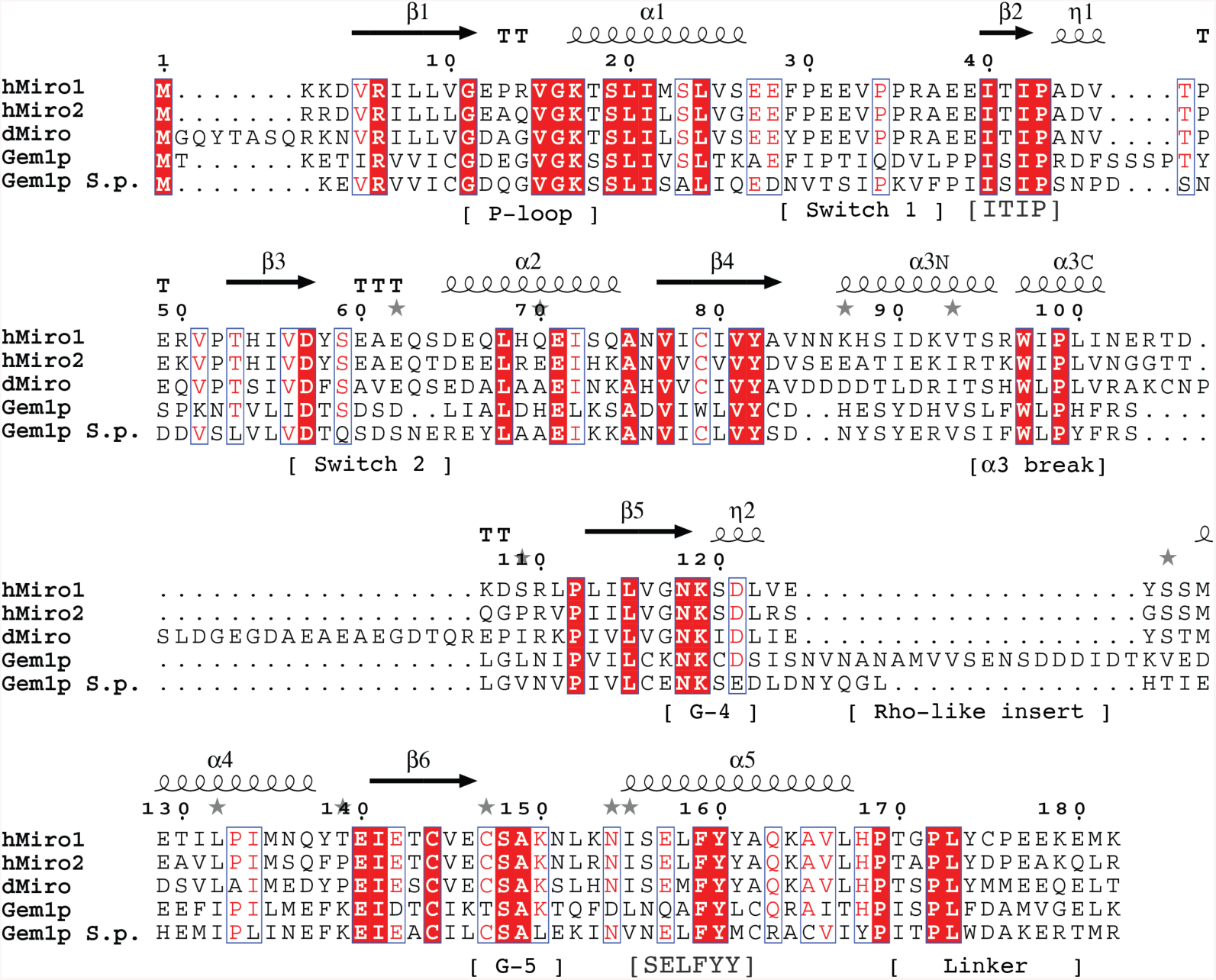
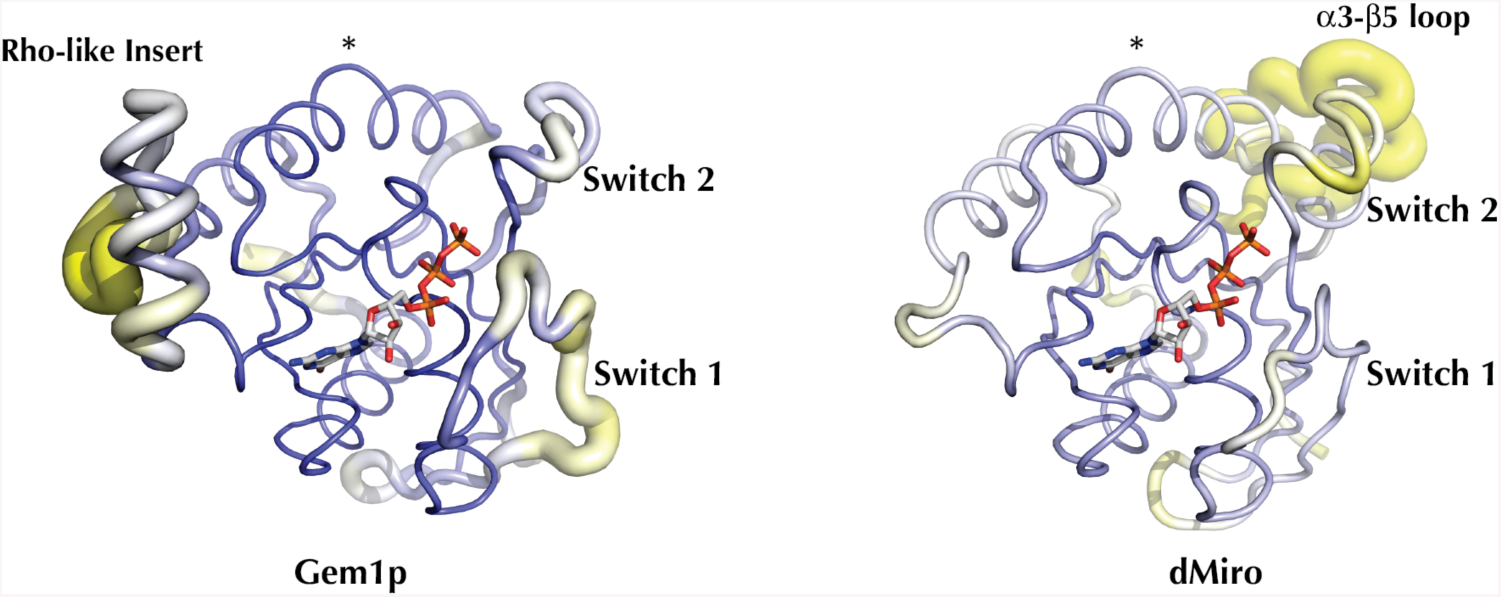
Conserved and divergent regions of the Miro N-terminal GTPases. (A) Sequence alignment of the nGTPase domains of HsMiro1/2 and Gem1p. The positions of the secondary structural elements are indicated above the alignment, and motifs referenced in the text indicated below the alignment. Residues 4-172 are visible in the crystal structure of the HsMiro1 nGTPase. Sequence alignment generated using T-Coffee and ESPrint (Di Tommaso et al., 2010; Robert and Gouet, 2014). (B) A homology model of Gem1p generated by I-Tasser (Zhang, 2009) using the HsMiro1 nGTPase structure as a template. The PyMOL ‘putty’ representation (Mura, 2004) encodes the estimated local accuracy of the model, and highlights that the ‘switch 1’ and ‘rho-like insert’ regions of Gem1p can not be modeled reliably due to divergence between HsMiro and Gem1p. The Miro and Gem1 structures deviate most significantly at Switch 1, and at the Rho-like insert. The position of the *α*3 break is highlighted with ‘*’. In contrast, the structure of the *Drosophila* DmMiro nGTPase (at right) can be modeled readily, with the exception of an extended loop inserted between *α*3 and *β*5 (labeled). In both images, the poorly ordered N- and C-terminal peptides are omitted – the Gem1p model includes residues 2 to 189 and the DmMiro model includes residues 8 to 198. The position of bound GTP is modeled based on alignment with the HsMiro1 GTP structure.

### 2.6. Size Exclusion Chromatography - Small Angle X-ray Scattering (SEC-SAXS)

SEC-SAXS Experiments were performed at the BioCAT beamline 18-ID-D at the Advanced Photon Source of Argonne National Laboratory (Mathew et al., 2004). A Superdex 200 Increase column was pre-equilibrated with 25 mM Tris, pH 8.5, 500 mM NaCl, 1 mM MgCl2, 0.5 mM TCEP, 2 μM GTP and either 1 mM CaCl2 or 1mM EGTA. Flow rate was 0.75 ml/min. Samples of HsMiro1 (residues 1-592 with a C-terminal 6xHis tag) and HsMiro2 (residues 1-588 with a C-terminal 6xHis tag), prepared in 25mM Tris pH 8.5, 500mM NaCl, 1mM MgCl2, 1mM CaCl2, 0.5 mM TCEP-HCl, and 2.5% sucrose, were injected onto the column (HsMiro1: 300 microliters at 2.2 mg/ml and 350 microliters at 5.8 mg/ml; HsMiro2: 400 microliters at 2.7 mg/ml). SAXS was performed in-line with this size exclusion column setup (Malaby et al., 2015). Photons scattered from the 12 keV X-ray beam were detected with a Dectris Pilatus 1M detector at a distance of 3.5 m from the sample. One second SAXS exposures of fractions of the SEC column, recorded every 3.0s, showed increased x-ray scattering at 3 distinct peaks, as shown in **Suppl. Fig. 3**. Only the major peak, which represents the monomeric species, was evaluated. Buffer background was obtained by radially binning each exposure, and then averaging over the first 100 fractions. The central portion of the monomer peak was averaged and background subtracted to obtain I vs. q curves (**Suppl. Fig. 4**, top panels) (Konarev et al., 2003). The radius of gyration Rg, zero-angle intensity I_0_, and associated uncertainties for these parameters were obtained by weighted linear regression of log(I) vs. q^2^ as shown in the Guinier plots in **Suppl. Fig. 4**. Dmax was determined from the pair distance distribution function P(r) with the program GNOM (Bussler et al., 1998; Svergun, 1992). In determining Dmax, GNOM gives a “total estimate”, which identifies common artifacts encountered by the GNOM method. Dummy atom modeling for SAXS reconstructions shown in **Fig. 3A** were done with the program DAMMIF (Franke and Svergun, 2009) (10 runs per SAXS structure) and aligned, averaged, and filtered using SUPCOMB (Kozin and Svergun, 2001), and DAMAVER/DAMFILT (Volkov and Svergun, 2003). SAXS parameters are reported in **Suppl. Table 2**. The A260/A280 ratios measured during the runs of 0.67 (HsMiro1 monomer) and 0.81 (HsMiro2 monomer) are consistent with theoretical values calculated from amino acid composition with two nucleotides bound (A260/A280 ratios of 0.70 and 0.75, respectively, when nucleotide-bound; 0.52 and 0.55, respectively, when nucleotide-free), however the nucleotide-bound state was not determined directly.

**Figure 3.**
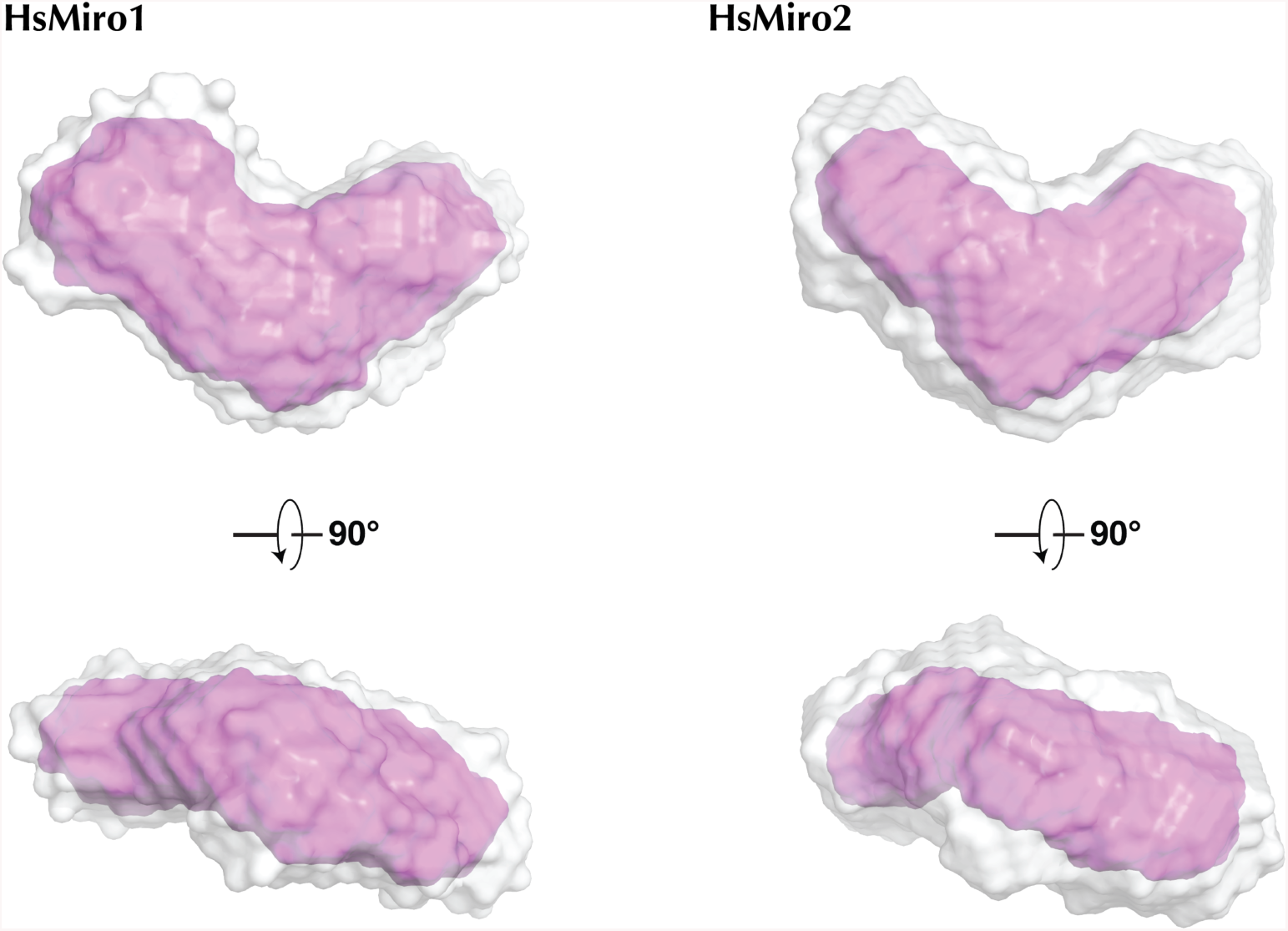
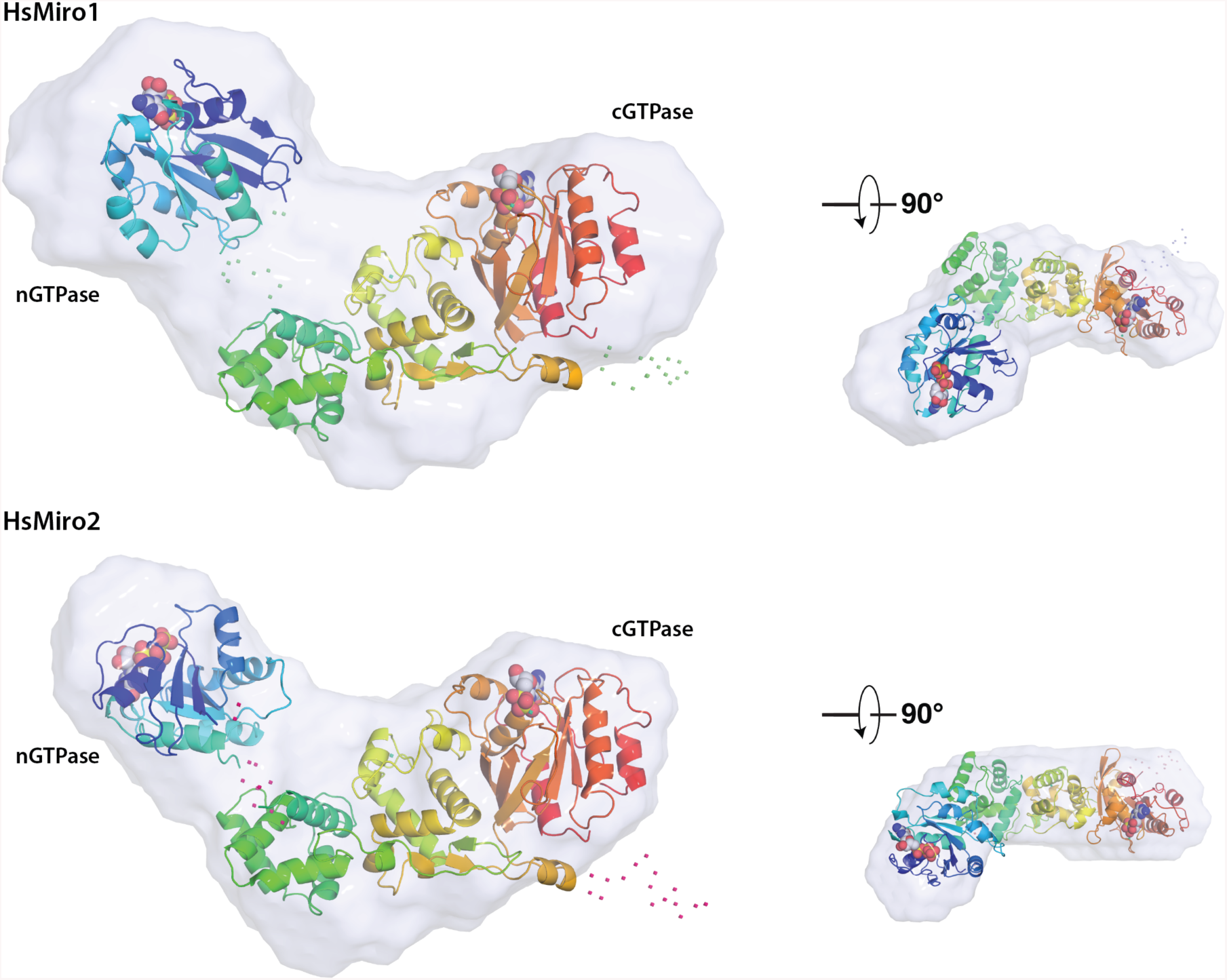
Solution structure of monomeric HsMiro1 and HsMiro2. (A) Averaged reconstructions from 10 DAMMIF calculations of the processed SAXS data from HsMiro1 and HsMiro2. The DAMAVER average of the DAMMIF runs is shown in gray and the DAMFILT filtered envelope is shown in pink (Franke and Svergun, 2009; Petoukhov et al., 2012). The dummy atoms used to construct the contours shown were generated with a radius of 4Å. (B) Representative monomer models generated using BUNCH (Petoukhov and Svergun, 2005) are overlaid with the corresponding SAXS scattering envelope reconstructions. The two models are aligned with respect to the MiroS domain, with the nGTPase at left in each case. The positions of the nucleotides bound to each of the GTPase domains are shown as spheres. Shown smaller at right are the overlays viewed after a ∼90° rotation around the horizontal axis. The scattering envelopes reasonably enclose both the rod-like MiroS and nGTPase domains. Note that while the orientations of MiroS in the two structures are similar, the orientations of the nGTPase domains are different, reflecting the fact that its position cannot be fixed based on this data; their relative dispositions suggest that the nGTPase domains are not tightly associated with the ELM1 domain of MiroS. Dots represent amino acids not observed in the crystal structures and modeled as flexible scattering elements.

### 2.7. SAXS Rigid-Body Modeling

A model of the full-length soluble domain of HsMiro was obtained using BUNCH (Petoukhov and Svergun, 2005). Rigid body optimization of theoretical scattering data to observed SAXS scattering data was based on partial scattering amplitudes obtained from the structures of the nGTPase of HsMiro1 and the EF-hand/cGTPase domains of HsMiro1 (PDB ID: 5KTY) subject to the constraints of the known linker peptide length between the two fragments. The program CRYSOL was used to generate theoretical scattering data from the nGTPase (residues 3-170, with the poorly ordered residues 171-175 deleted) and MiroS (residues 181-582) structures. The Pre-BUNCH input specified the full sequence of the soluble domain (1-600), such that contributions of three peptides (1-2, 171-180, 583-600) were modeled as unknown. Default parameters were used throughout. The modeling runs were carried out against the observed SAXS scattering data obtained from both HsMiro1 and HsMiro2, yielding χ^2^ values of 1.137 and 4.137 for the two HsMiro1 samples, respectively, and a χ^2^ of 1.007 for HsMiro2. The generated models of HsMiro1 and HsMiro2 obtained by rigid body optimization against the scattering data were then superimposed on the DAMMIF dummy atom reconstruction obtained from each dataset using SUPCOMB (Kozin and Svergun, 2001). The normalized spatial discrepancies (NSD) obtained following overlap of the protein model with each dummy atom reconstruction were: HsMiro1 2.0055 and 2.1277; and HsMiro2 1.7645. These superpositions for HsMiro1 and HsMiro2 are shown in **Fig. 3B**.

## 3. Results and Discussion

The crystal structure of the Human Miro1 N-terminal GTPase was determined at 1.72 Å resolution with magnesium and GTP bound (**Table 1**), with two molecules in the asymmetric unit. Both polypeptide chains are bound to Mg^2+^GTP and are very well defined in the electron density map. The overall fold of the HsMiro1 nGTPase is similar to that of other small GTPases (**Fig. 1B**). The structure can be superimposed on the structure of Ras-GTP with an RMS deviation of 0.902 Å (over 99 C*α* atoms, PDB: 5P21), and on the structure of Rho-GTP with an RMSD of 0.700 Å (over 107 C*α* atoms, PDB: 3TVD). Portions of the C-terminal peptides of the two protomers, residues 169-172/175, interact across the dimer interface but are poorly ordered, consistent with their likely role as a flexible ∼10 residue linker between the N-terminal GTPase and the remainder of the protein, MiroS (Klosowiak et al., 2013). The overall structure is shown in **Figure 1B**.

### 3.1. Binding interactions include unusual coordination of Mg^2+^ ion

Small GTPase binding sites are generally characterized by five conserved sequence motifs (G-1 to G-5) (Bourne et al., 1991). In the HsMiro1 nGTPase, the mainchain atom hydrogen-bonding interactions between the ‘P-loop’ (G-1) motif and the phosphate groups of the bound GTP are typical of members of the small GTPase family. The P-loop also contributes two sidechains, Pro13 and Arg14, which may serve to sequester access to the active site. Both the Switch 1 (G-2) and Switch 2 (G-3) motifs, which in other GTPases mediate catalytic activity, appear to be well-ordered and show extensive interactions to residues of the active site (**Suppl. Fig. 1B**). However, as the structure contains bound GTP, the configuration of the active site residues and the accompanying water structure seen in the crystal structure are apparently non-catalytic. Interestingly, the magnesium ion coordination mediated by the ‘Switch 1’ (G-2) motif is contributed not by a sidechain atom (e.g. a Thr side chain hydroxyl as is seen in Ras (Bourne et al., 1991)), but rather by the carbonyl oxygen of Pro35 (**Fig. 1C** and see **Suppl. Fig. 2**), which is one of a proline-proline (PP) sequence pair the first residue of which is highly conserved among the Miro GTPases (**Fig. 2A**). There is no threonine sidechain in the vicinity, obviating canonical Switch 1 Thr coordination that would be inferred by analogy to Ras (Frederick et al., 2004; Kornmann et al., 2011). In the yeast Gem1p, to the extent that the Gem1p Switch 1 region can be modeled (see below), a threonine residue (Thr33) occurs six residues N-terminal to the position of the PP motif; its mutation to alanine was found to have no effect on subcellular localization (Kornmann et al., 2011; Frederick et al., 2004). This direct mainchain coordination of magnesium is unusual in small GTPases but not unprecedented, as similar carbonyl coordination is found in structures of the RhoA GTPase GDP complex (Wei et al., 1997; Pellegrini and Bowler, 2016). In the nGTPase, the G-3 (Switch 2) motif contributes three sidechains that are directed towards the active site – Asp57, which canonically stabilizes Mg^2+^ coordination, Ser59, which appears to form a labile hydrogen bonding interaction with an active site water, and Ala61, which forms a ‘cap’ at one side of the active water structure (**Fig. 1C**). The water structure is well-defined, with minor differences between the two protomers within the asymmetric unit. In both sites, a water molecule bridges an oxygen of the GTP γ-phosphate with the backbone amide of Gln60, with two additional water molecules completing a tetragonal hydrogen bonding arrangement. This arrangement is conserved, with the addition in one site of the asymmetric unit of an adjacent water molecule bridging and possibly stabilized by both the sidechain hydroxyl of Ser59 and γ-phosphate oxygens of GTP (**Fig. 1C**). While a water at this position might play a functional role at the active center of the GTPase, how these interactions might have significance here remains unknown. Finally, the interactions of the G-4 motif with the guanine base are mediated canonically by the sidechains of Asp121 (hydrogen bonding to the purine) and Lys 119 (packing against it). The exocyclic oxygen of GTP buried within the active site is stabilized via a hydrogen bond to the mainchain nitrogen of Ala149, which is part of the conserved G-5 motif that occurs just prior to the C-terminal helix of the domain.

### 3.2. GTP binds tightly to the crystallized N-terminal GTPase domain

The HsMiro1 nGTPase was initially crystallized in the presence of a mixture of 1mM GDP and GTP (see Methods), however, the bound ligand evident in the resulting 3.2A electron density map could be readily modeled as a molecule of GTP. Despite the presence of GDP during crystallization, binding heterogeneity could be excluded both by the quality of the density and by the consistency of the B-factors across the phosphate chain with GTP modeled into the site. Subsequently a 1.7Å resolution dataset was obtained from a crystallization carried out in the presence of only a low concentration (1μM) of added nucleotide, and this structure, reported here, also revealed well-defined electron density consistent with binding of only Mg^2+^-GTP within the active site (**Fig. 1B**). We conclude, therefore, that GTP is very tightly bound to our HsMiro1 nGTPase domain construct and believe that the bound species originated from nucleotide endogenous to the expression system that co-purified with the protein.

However, the co-purification and evident stability of the GTP observed bound in our crystal structure is surprising, as previous work reported GTP hydrolysis by the purified *Drosophila* Miro (Lee et al., 2016), provided qualitative evidence for hydrolysis by a human nGTPase fusion (Suzuki et al., 2014), and found robust enzymatic activity with the purified yeast homolog Gem1p (Koshiba et al., 2011). More recently, it has been shown that the HsMiro nGTPase itself can exhibit readily measurable GTP hydrolysis activity (Peters et al., 2018). The expression constructs used in the latter study comprised residues 2-169 of HsMiro1/2, terminating just C-terminal to helix α5, and are quite similar to the construct that was crystallized, residues 1-180 of HsMiro1. We cannot yet reconcile our observations with these data, although we note that the additional C-terminal 11 residues are poorly ordered in our structure, and are adjacent to a conserved interface that mediates the non-crystallographic dimer (discussed below). We speculate that the difference in hydrolysis behavior might arise from differences in the protein constructs and purification methods used, but further study will be required to resolve this issue.

### 3.3. Structural relationships to the Rho and Miro C-terminal GTPases

Based on a number of sequence divergences Miro is no longer considered a *bonafide* member of the Rho GTPase family (Koshiba et al., 2011; Wennerberg and Der, 2004). Most notable is the absence of the so-called ‘rho-helix’, a conserved ∼13 amino acid sequence inserted between the G-4 motif and helix α4 of the Ras-like GTPase fold (Wennerberg and Der, 2004). Instead, in HsMiro seventeen residues of the rho-helix loop are replaced by a much shorter 4-residue turn. Nevertheless, several elements characteristic of the Rho GTPases are conserved between Miro and Rho and are structurally superimposable: First, a highly conserved Phe/Pro motif N-terminal to the ‘Switch 1’ region packs against the guanine base. Second, a highly conserved WxP motif in helix α-3 serves to maintain the structural relationship between helices α1, α2, and α3 by introducing a kink in helix α3 at the same position in almost all rho GTPases (**Fig 1B**). Finally, the G-5 motif sequence CSAK characteristic of rho GTPases is structurally conserved, as the serine and alanine residues both play a role in positioning the guanine base. Interestingly, while Pro35 of the PP motif (above) of Switch 1 appears to be highly conserved in Rho GTPases, the structure of this region, termed the ‘core effector domain’ (Wennerberg and Der, 2004), is generally quite different in Rho GTPases from that in HsMiro1, and the position and orientation of the ‘conserved’ proline is distinct. The absence of several other sequence elements conserved in the Rho family, particularly large hydrophobics that appear to function to position the core effector domain in rho GTPases, suggests that the apparent conservation of the proline here does not reflect similarity of structure or function.

The recent identification of the cGTPase as an NTP hydrolase (Peters et al., 2018) suggests that the two GTPase domains of Miro are mechanistically different, and, indeed, they are structurally quite distinct (Klosowiak et al., 2016) with an overall RMSD for their superposition of 2.09 Å over 104 C*α* atoms. The nGTPase domain of Miro exhibits only 22% sequence identity with its cGTPase domain, and notably, sequence and structural alignments show almost no similarity in the switch regions between the nGTPase and cGTPase (**Suppl. Fig. 1**). Curiously, the sequences of the Miro cGTPase Switch regions themselves are poorly conserved within the Miro protein family (**Suppl. Fig. 1A**). In contrast to the nGTPase, the Miro cGTPase domain readily exchanges nucleotide, and its structures in GDP-, GDP*Pi-, and GMPPCP-bound states have been determined (Klosowiak et al., 2013; 2016). Particularly notable is the location of its Switch 1 region (**Suppl. Fig. 1B**), which adopts an open conformation distant from bound nucleotide. The latter is consistent with the possibility that its catalytic and regulatory mechanisms may be unrelated to that of the broader family of GTPases.

### 3.4. Structural dissimilarities between the HsMiro and yeast Gem1p nGTPases

The structure of the N-terminal domain of the HsMiro1 potentially provides a structural model for the yeast homolog Gem1p. However, while the overall sequence identity between human and yeast is 30% over the nGTPase of Miro, there are two interesting features that suggest that structural and functional inferences may be problematic - which may be particularly important given the apparently distinct functional roles played by the homologs in yeast and metazoans. It has previously been noted that Gem1p nGTPase lacks both a rho-like ‘Switch 2’ (G-3) sequence consensus and rho-like helix sequence (Frederick et al., 2004). In contrast to HsMiro, however, a ∼17 residue insertion occurs in Gem1p at the position of the ‘rho helix’ (**Fig. 2B**), which though divergent in sequence, retains the polar character (Wei et al., 1997) of the rho-specific insert (9 of 17 residues are Asn/Glu/Asp). This sequence is absent in HsMiro1 and hence we cannot model it. There is evidence for conservation of the sequence of the G-3 (switch 2) motif amongst Miro nGTPases (**Fig. 2A**), most notably Ser59 which is directed towards the water structure at the putative active center in HsMiro1. Interestingly, however, the G-2 (switch 1) motif is poorly conserved - although a ‘PP’ motif is present in the yeast sequence, its position relative to elements that can be reliably modeled based on our structure (i.e. which correspond to the core beta strands of the nGTPase fold) is shifted four residues C-terminal; consequently it is impossible to model reliably the structure of this region, as shown, graphically, in **Fig 2B**. Since both the G-2 and G-3 GTPase motifs play a direct role in GTP hydrolysis and exchange (Cherfils and Zeghouf, 2013), differences in their structural arrangements and nucleotide interactions presumably lead to distinct functional and catalytic behaviors. The extensive interactions of the G-2 motif in the HsMiro nGTPase domain with nucleotide may significantly stabilize the bound species, consistent with our observation of co-purification of GTP. If those interactions were absent or different in the yeast homolog, the biochemical behavior of the two proteins would likely be quite distinct.

### 3.5. HsMiro1/2 is an elongated ‘crescent’ in solution

With structures of both the HsMiro1 nGTPase and the HsMiro1 ‘S’ domain comprising the EF-hand and cGTPase domains (Klosowiak et al., 2016) available, we sought to study the assembly of the full-length HsMiro with the goal of obtaining a model of the structure of the complete extramembranous Miro polypeptide. Our previous SAXS studies of the DmMiroS (Klosowiak et al., 2013) indicated good agreement with the structure determined crystallographically. However, in those studies of the *Drosophila* Miro we noted some heterogeneity in the SAXS data from the full-length protein likely arising from aggregation and/or formation of multimers. To obviate this previously noted heterogeneity, we took advantage of a SEC-SAXS facility available at the BioCAT beamline that allowed us to readily resolve scattering from dimers and aggregates from that of the monomeric full length protein (Malaby et al., 2015) (**Suppl. Fig. 3**). Experiments were carried out with two independently purified samples of full-length HsMiro1, and from a sample of its paralog HsMiro2. Measurements carried out in the presence and absence of 1 mM calcium chloride revealed a negligible effect on the SAXS scattering parameters for HsMiro1/2, consistent with our previous studies showing that changes in calcium occupancy of the EF hands of MiroS altered neither the solution structure nor the crystal structure of MiroS (Klosowiak et al., 2013). Significantly, the scattering parameters were remarkably consistent among the three samples (**Suppl. Fig. 4**). We generated *ab initio* scattering models from each dataset. The reconstructions revealed an overall crescent-shaped structure which was similar amongst the HsMiro1 and HsMiro2 samples (**Fig. 3A**), and which was consistent with the rod-like shape of the fragment of HsMiro determined previously (Klosowiak et al., 2013; 2016). The reproducibility of the reconstructions obtained was indicated by DAMMIF normalized spatial discrepancy (NSD) values of 0.62 and 0.63 for HsMiro1 and HsMiro2, respectively (**Suppl. Table 2**). From this we conclude that each of the reconstructions generated for HsMiro1/2 are both similar to each other and, as expected from our crystal structures, do not exhibit substantial inter-domain disorder.

We generated structural models of the full-length Miro1/2 using the programs BUNCH (Petoukhov and Svergun, 2005) and CORAL (Petoukhov et al., 2012), combining our SAXS data with the previously reported crystal structures of the HsMiro1 EF-hand and cGTPase domains (‘MiroS’; PDB ID: 5KTY) and the nGTPase domain. We included the unmodeled N- and C-termini, and modeled the short linker peptide between the nGTPase and MiroS (170-180) as a flexible restraint between two rigid-body domains. The overall fit of the resulting structural models yielded in the best case *χ*^2^ values of 1.137 for HsMiro1 and 1.007 for HsMiro2. In each of the models we found the position of the nGTPase domain to be consistent, located in one lobe of the ‘crescent’ but extended from the remainder of the MiroS (**Fig. 3B**). While the position of the nGTPase relative to MiroS must be constrained by the relatively short linker between the two, its positions in our models suggests that its relative orientation is less so, and that the nGTPase does not form a stable interface with the rest of the molecule. Thus, the deviation of the *χ*^2^ values may arise both from the poorly defined polypeptide ‘linker regions’ (at the N- and C-termini, and the nGTPase boundary, and internal to the otherwise rodlike assembly of the HsMiro S domain), as well as from the distribution of orientations available to the nGTPase relative to the rest of the protein. Nevertheless, the consistent observation of a similar overall shape derived from the scattering data obtained from the three samples is striking. We conclude that the nGTPase domain has limited mobility relative to the remainder of the molecule, but, in contrast to the extensive interface of the cGTPase domain the second ELM2 domain (Klosowiak et al., 2013), the nGTPase domain of HsMiro does not appear to be tightly coupled to its EF-hand domains. We speculate that such conformational freedom could facilitate interactions with distinct binding partners across multiple interfaces of the nGTPase domain.

### 3.6. A conserved motif mediates a potential binding interface

The HsMiro1 nGTPase construct crystallized here appears to be monomeric in solution (Klosowiak et al., 2016). However, in the crystal structure, the nGTPase domain associates as a two-fold symmetric non-crystallographic dimer **(Fig. 4A)**. The interface between the two protomers is extensive, with a buried surface area of 1225 A^2^, and a computed *Δ*G for dimer formation (Krissinel and Henrick, 2007) of −9.1 kcal/mol. The interface involves extensive interactions between helices α4, the bridging β-strand, and helix α5, and remarkably, it is mediated by previously unrecognized but highly conserved sequence motif of the Miro nGTPases (**Fig. 4B**), which we term ‘SELFYY’. The tyrosines of the motif occur along the C-terminal helix, α5, of the nGTPase fold, and they contribute to formation of a complex surface that involves numerous hydrophobic and polar interactions. The tyrosines of the motif are absent in other Rho GTPases, and indeed the fingerprint of the interface rationalizes much of the Miro family sequence conservation observed throughout this region of the molecule **(Fig. 4C)**. Consequently, we infer that the dimer interface observed in the crystal reflects a functionally important interaction of the Miro nGTPase domain. Of note is that Ser156 (‘S’ of the SELFYY motif) is phosphorylated by PINK1 kinase thereby promoting recruitment of the ubiquitin ligase Parkin and triggering Miro ubiquitination and degradation (Shlevkov et al., 2016; Wang et al., 2011).

**Figure 4.**
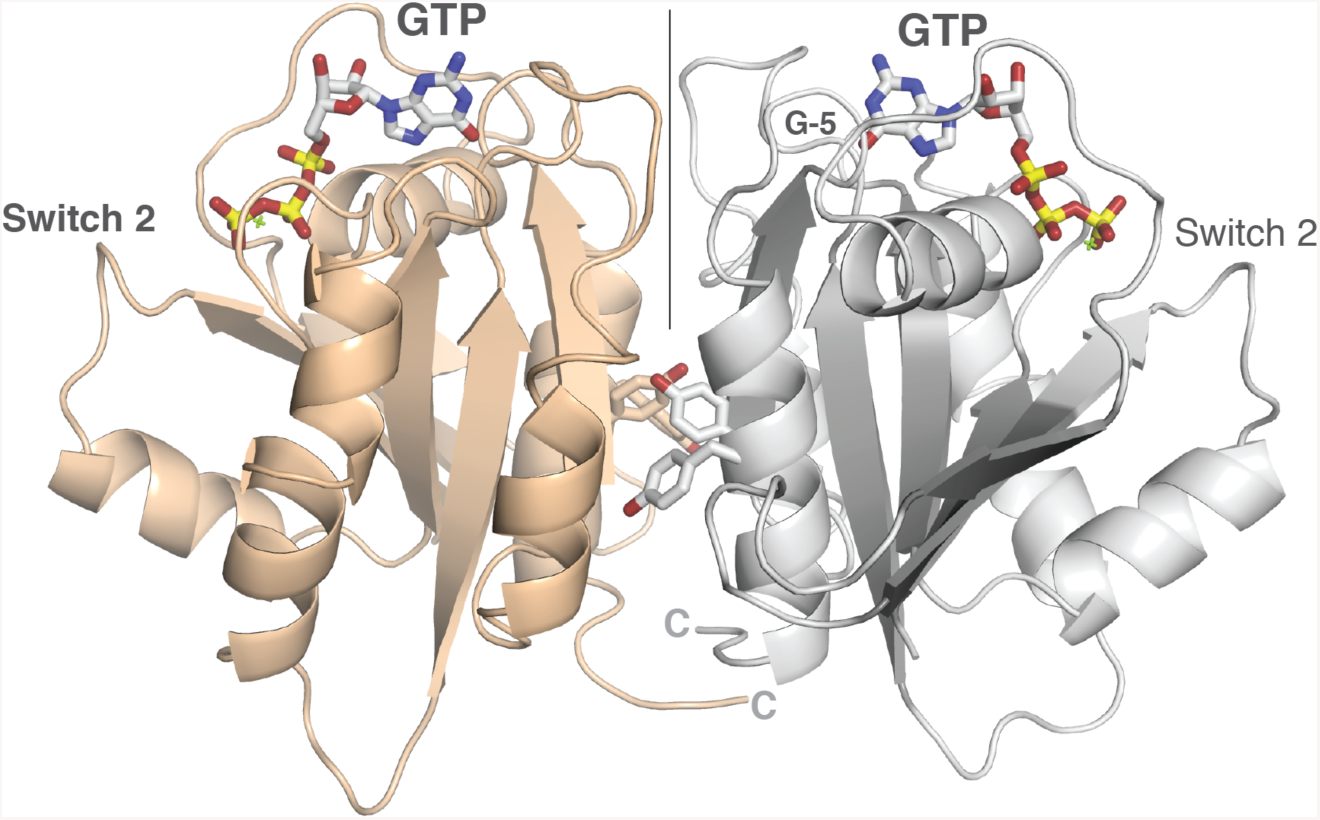
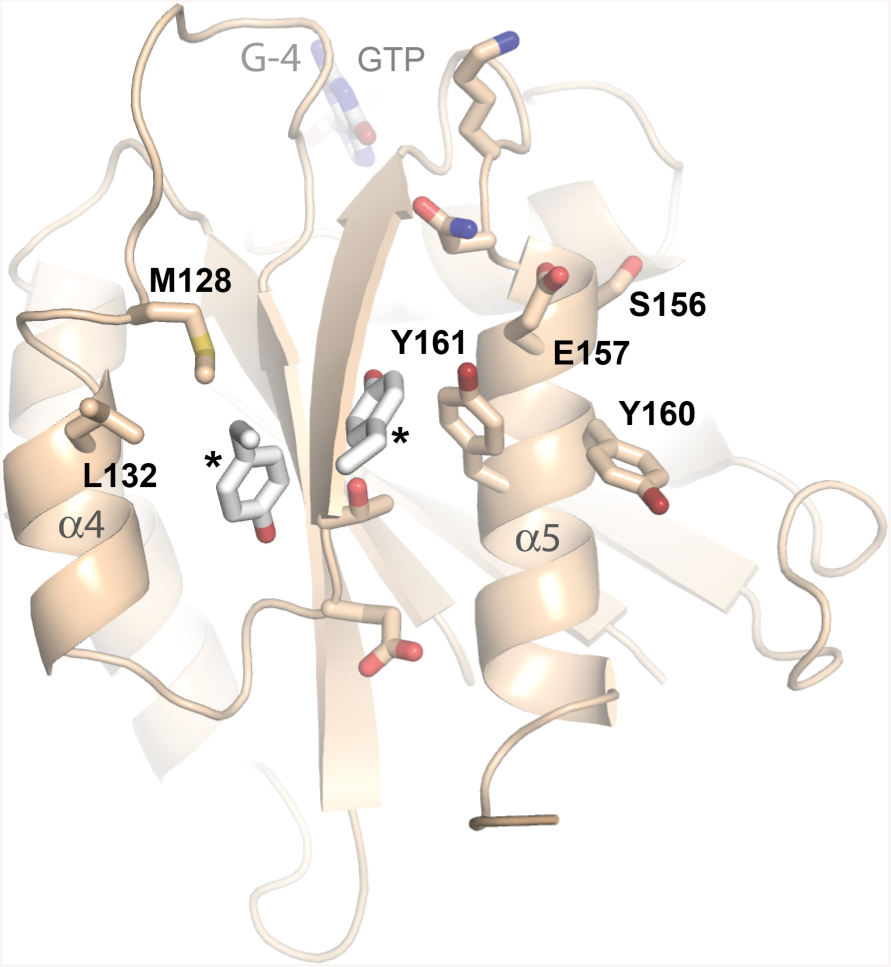
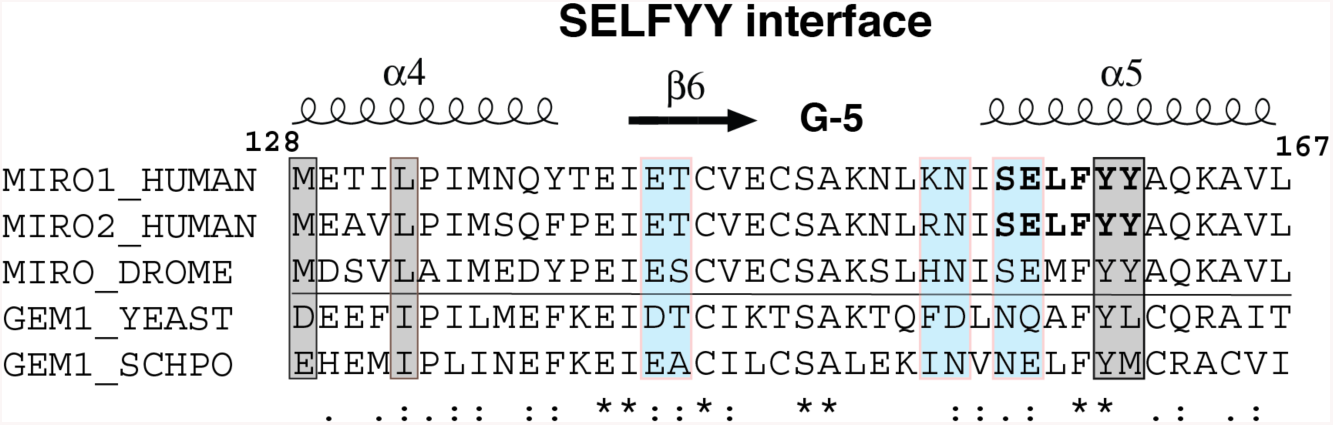
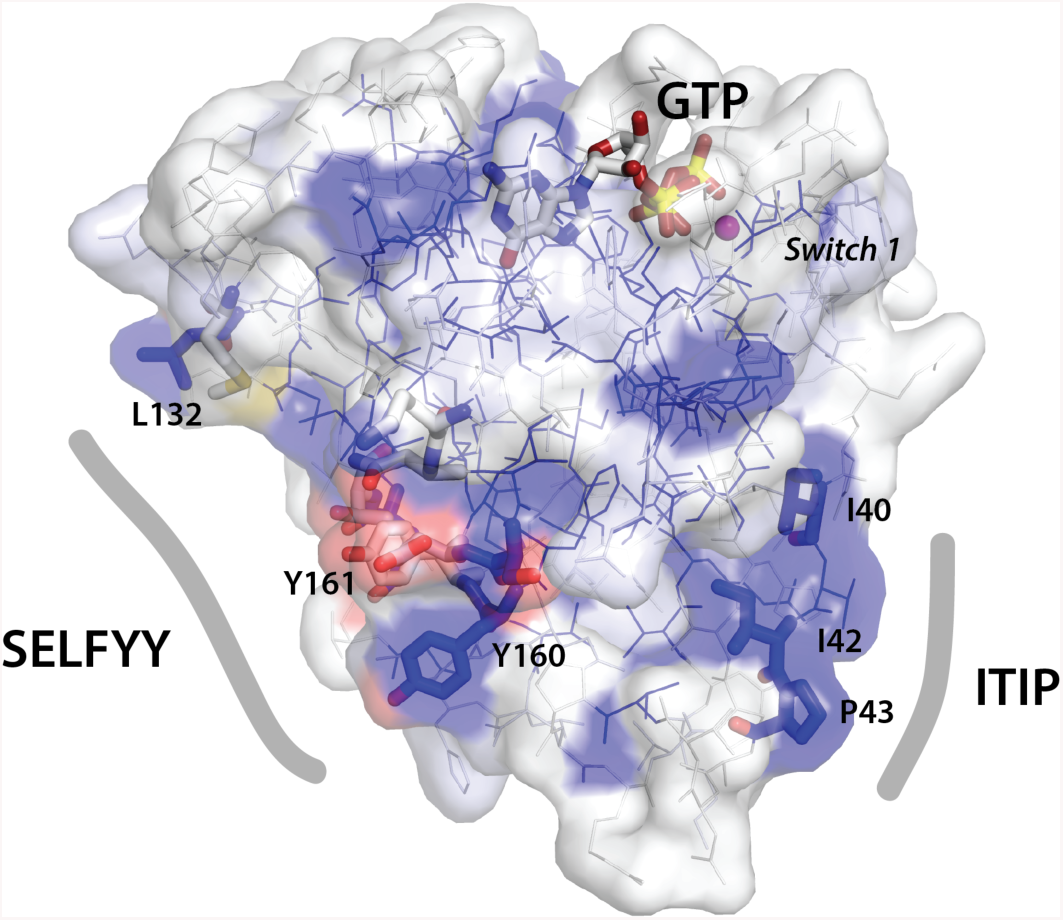
Dimerization and conserved surfaces of the HsMiro nGTPase. (A) Dimerization of the nGTPase domain in the crystal. The protomers associate across a non-crystallographic two-fold axis. At the center of the interface is a cluster of paired tyrosine sidechains of the ‘SELFYY’ motif. The positions of the GTP molecules are indicated. Note that the interface is distal to the Switch 1 and Switch 2 regions, and therefore is unlikely to be responsive to a specific nucleotide binding state. The G-5 motif in the foreground at right is indicated. (B) Details of the SELFYY motif interface. One monomer of the dimer is shown rotated 90° relative to (A), with residues substantially buried at the interface shown as sticks. The hydrophobic residues of the interface (M128, L130, Y160, Y161) are labeled, with the positions of the sidechains of the tyrosines extending across the dimer interface (‘*’) shown to highlight their intercalation into a pocket formed by sidechains extending from the α4 and α5 helices. The G-4 loop and GTP are indicated. S156 has been reported to be phosphorylated by PINK1 kinase (Shlevkov et al., 2016). (C) Sequence conservation in the SELFYY interface of the Miro nGTPases. The surface residues that are substantially buried upon formation of the interface are highlighted – hydrophobics in grey, polar residues in teal. The associated secondary structure is indicated above the alignment, sequence conservation indicated below it. (D) Surface representation of the nGTPase domain. Sequence conservation is mapped as surface color from white (not conserved) to blue (highly conserved). The sidechains of the SELFYY interface and the conserved ‘ITIP’ hydrophobic surface are shown as sticks, and the SW1 region is indicated. The GTP binding site is at top, with GTP shown as sticks. At left, the ‘SELFYY’ surface extends across a broad face of the domain (L132 – Y160); at right the ‘ITIP’ hydrophobic surface (I40 – P43) follows switch 1 (‘SW1’).

The position of the SELFYY interface along a face of the protein fold distal to the switch regions of the nucleotide-binding site suggests that the interface is not responsive to nucleotide-binding state. However, interestingly, a survey of crystal structures of H-Ras has shown that a dimeric form mediated by the corresponding interface, termed the α4/α5 dimer, is preferentially observed when Ras is in its ‘active’ or GTP-bound state (Spencer-Smith et al., 2017; Güldenhaupt et al., 2012). This region of the Ras protein has been termed the ‘allosteric lobe’, and although the functional significance of dimerization in Ras is somewhat controversial, there is evidence that it may be functionally coupled to its activity (Spencer-Smith et al., 2017; Fetics et al., 2015). These results raise the interesting, albeit speculative, possibility that the unexpected stabilization of GTP observed in our crystal structure might be coupled to formation of the SELFYY-mediated dimer.

There is an additional conserved hydrophobic surface of the nGTPase, which we term ‘ITIP’ (**Fig. 2A**), that contributes to a different crystal contact, burying 599 A^2^ with a calculated *Δ*G of −7.8 kcal/mol, as evaluated using PISA (Krissinel and Henrick, 2007). The motif is surface exposed and is situated adjacent to the switch 1 region of the nGTPase (**Fig. 4D**). The conservation of these two sequence motifs in Miro suggests that they may both be functionally important. However, the low resolution of SAXS data and the multiple modeled orientations consistent with it do not allow us to establish the relative orientations of the ‘ITIP’ or ‘SELFYY’ motifs of the nGTPase with respect to the HsMiroS domain. The sterically accessible orientations of the nGTPase modeled within the scattering envelope suggest that one or both might mediate association of the nGTPase with the rest of the intact Miro. However, in the BUNCH models, the SELFYY motif in particular is potentially exposed at the surface of the HsMiro1/2 extra-membranous domain (**Fig. 3B**). HsMiro1 has been shown to occur in a protein complex with HsMiro2 (Kanfer et al., 2015; van Spronsen et al., 2013; Weihofen et al., 2009), and its binding partners Kinesin-1, TRAK1/2, DISC1, Myosin19, OGT, Mfn1/2, p150glued, and PINK1 self-associate in cells (Koutsopoulos et al., 2010; Weihofen et al., 2009; Cao et al., 2017; Leliveld et al., 2009). We speculate, therefore, that one or both of these putative binding surfaces of the HsMiro1 nGTPase domain could mediate dimerization of Miro and/or functionally interact with its binding partners.

## 4. Conclusions

Here, we report the structure of the nGTPase of HsMiro1 in complex with GTP. We found no evidence for hydrolysis of GTP and conclude that the nucleotide is tightly bound and that the crystallized N-terminal GTPase construct is catalytically inactive. Extensive interactions with the ligand, including a novel backbone carbonyl coordination of magnesium, appear to stabilize the bound Mg^2+^GTP. The absence of hydrolysis is unexpected, as recent studies using a similar nGTPase construct have shown GTP hydrolysis activity similar to that of the small GTPase Ras (Peters et al., 2018). Additionally, Miro GTP hydrolysis activity has been measured in both the yeast Miro ortholog Gem1p, and the *Drosophila* DmMiro (Koshiba et al., 2011; Lee et al., 2016). Mutational studies have established that disruption of the putative active centers of both the N- and C-terminal Miro GTPases have phenotypic effect (MacAskill et al., 2009a; Murley et al., 2013; Saotome et al., 2008), however it remains to be established whether these mutations stabilize conformationally distinct ‘GTP-bound’ and ‘GDP-bound’-like states (as has been claimed (Fransson et al., 2003)), or whether they more simply disrupt nucleotide binding interactions thereby destabilizing the GTPase domain structure and/or its functional interactions. Further studies addressing the regulation of the GTPase activity of the nGTPase domain, whether by intrinsic and/or extrinsic interactions, will be needed to resolve these issues.

We also report a low resolution envelope for the structure of the intact HsMiro1 and HsMiro2 based on small angle X-ray scattering (SAXS) data, revealing a crescent-shaped structure consistent with positioning of the nGTPase domain at one end of, but not colinear with, the rod-like MiroS (comprising the two EF-hand and cGTPase domains). The short linker peptide between the nGTPase and the rest of the protein (∼10 residues) likely constrains its position. However, the low resolution of the scattering envelope does not allow determination of the extent to which the relative positions of the nGTPase and MiroS domains under these solution conditions are fixed or conformationally dynamic. The structure does seem to exclude direct interaction of the nGTPase with the face of the ELM1 domain (Klosowiak et al., 2013).

Miro has been found in the context of the cell to occur as a protein complex (Glater et al., 2006), and many of its reported binding partners are thought to be dimeric or multimeric (e.g. Kinesin-1, TRAK1/2, Mfn1/2) (Shlevkov et al., 2016; Lovas and Wang, 2013; Cao et al., 2017; Daumke and Roux, 2017). The exact stoichiometry and oligomeric state of Miro in its complexes *in vivo* is unknown. Here, we locate two conserved and largely hydrophobic surfaces of the nGTPase domain that potentially function as binding interfaces. The first, which we term ‘SELFYY’ is quite extensive and mediates homodimerization in the crystal, suggesting the intriguing possibility that situated at the constrained surface of the mitochondrial membrane it might mediate dimerization of the intact Miro as well. Based on its relationship to the GTPase protein fold, this surface is less likely to be responsive to nucleotide-binding state than the canonical ‘switch’ regions that are located adjacent to the phosphate chain. We also identify a second interface in the crystal that is mediated by conserved residues adjacent to the nGTPase ‘switch 1’ region. However, even if functional as *bonafide* interaction motifs, their roles are likely regulated by the presence of ligands and/or macromolecular binding partners. Further studies will be needed to determine whether and how each of these interfaces contributes to Miro function.

The paralogs Miro1 and Miro2 are differentially expressed in cells (Fransson et al., 2003) and it is becoming clear that their roles are functionally distinct (López-Doménech et al., 2018). Studies of Miro1 in mouse models have shown that Miro2 protein levels are not sufficient to replace Miro1 knockout dysfunction at a subcellular, cellular or organismal level (Nguyen et al., 2014; López-Doménech et al., 2016). Differential binding localization studies suggest that Miro1 and Miro2 interact differently with the motor adapters TRAK1 and TRAK2 (van Spronsen et al., 2013). The nGTPase domains of HsMiro1 and HsMiro2, which are believed to play the key role in the interactions with TRAK1/2 (Bocanegra et al., 2019; Oeding et al., 2018), are structurally quite similar, as they exhibit 73% sequence identity and 88% sequence similarity. Consequently, the set of 21 nonhomologous amino acid substitutions between them presumably differentiate the functions of the two proteins. Mapping these differences on the structure of HsMiro1 we find, interestingly, that they locate to one face of the domain, primarily along loops adjacent to the GTP binding site and to the C-terminal end of the switch 2 helix (**Fig. 5**). Notably, the ‘SELFYY’ surface, opposite, is highly conserved between HsMiro1 and HsMiro2, perhaps consistent with a common function as an interaction interface.

**Figure 5.**
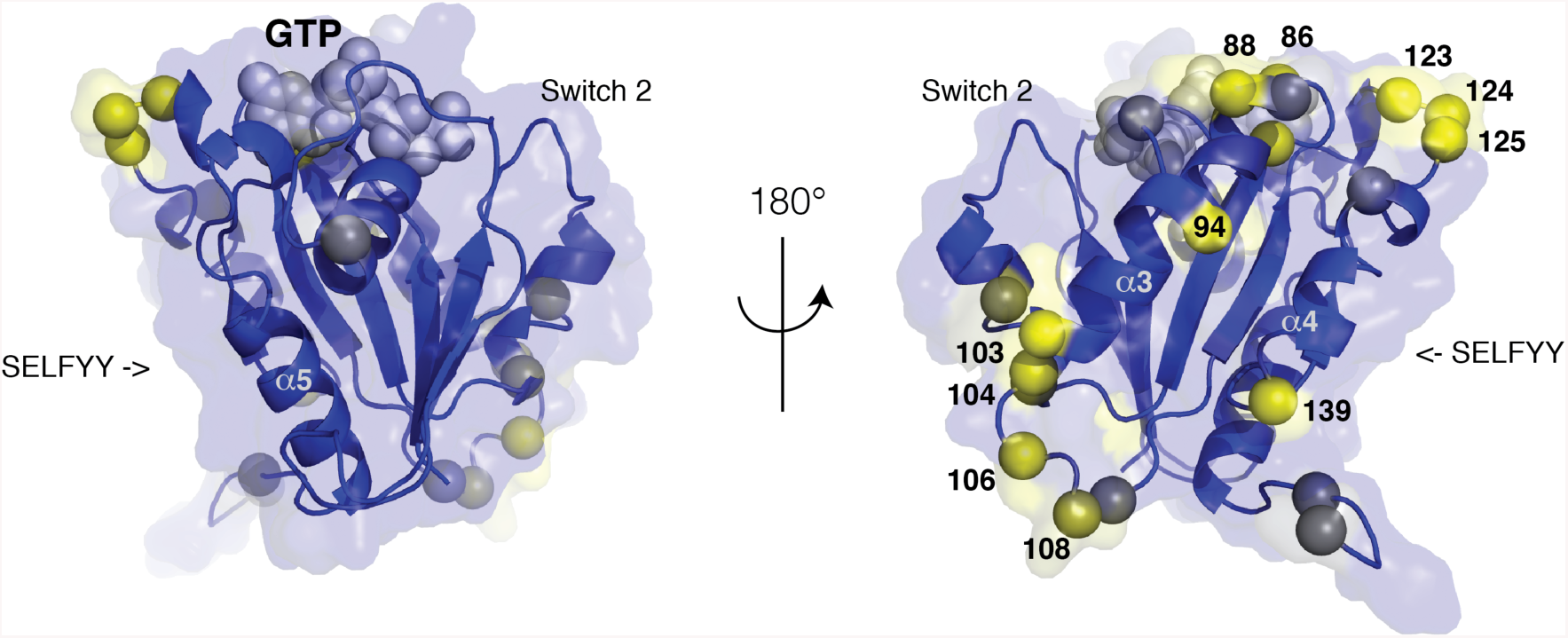
Differences between the nGTPases of HsMiro1 and HsMiro2 are localized to one face of the domain. The sequences of the nGTPases of HsMiro1/2 are 73% identical and 88% similar, with no gaps. The location of the 21 non-homologous substitutions are mapped on the structure of HsMiro1, using a color scale from blue (conservative) to yellow (non-conservative substitution). The two orientations related by 180° highlight the sidedness of the distribution of sequence differences. At right, towards the helices α3 and α4 of the GTPase fold, clusters of sequence divergence occur; select amino acid positions are labeled. Both views are perpendicular to the SELFFY interface, which is highly conserved. The positions of the Switch 1 and Switch 2 loops are indicated, and a CPK model of the GTP nucleotide is shown in grey.

Miro presents an interesting puzzle in that it is a calcium-regulated protein that contains two distinct GTPase domains of unknown function. Previous studies of HsMiro GTPase function primarily focused on observation of mitochondrial dynamics phenotypes in cells (MacAskill et al., 2009a; Murley et al., 2013; Saotome et al., 2008), and studies of the GTPase domains have been largely framed in terms of canonical models for GTPase structure and function (Bourne et al., 1990; 1991). Whether the Miro GTPases even act, like Ras, as ‘on/off switches’, or whether they serve assembly-activated ‘docking’ modules (Focia et al., 2004; Qi et al., 2016), remains unknown. By providing a structural model of the complete extramembranous HsMiro polypeptide and identifying surfaces that are likely to be functionally important, our structures should provide a framework for more extensive interrogation of the structure/function relationships of this important mitochondrial protein.

## Author Contributions

S.E.R. and D.M.F. supervised the research. K.P.S. designed and performed all experiments. K.P.S. purified all proteins with assistance from J.L.K. K.P.S. and P.J.F. collected and processed X-ray diffraction data. K.P.S. solved the phases. D.M.F. and K.P.S. interpreted the structures. K.P.S., D.M.F. and S.C. collected, processed, and analyzed SEC-SAXS data. E.C.L assisted in interpretation of the SAXS data. J.L.K initiated crystallography to the project and contributed original ideas. K.P.S and D.M.F wrote the manuscript with assistance and input from all authors.

## Acknowledgements

We thank Samuel Light and the Anderson lab for crystallization supplies and thoughtful feedback. We thank Melissa Gonzalez for assistance, and thank Theint Aung, Chi-Hao Luan, and David Haselbach for assisting with data collection and interpretation. We thank Priscilla Yeung and Jodi Nunnari for thoughtful discussions. K.P.S. and J.L.K. were supported by T32GM008382 and the Walter S. and Lucienne Driskill Graduate Training Program in Life Sciences at Northwestern University. S.E.R. was supported by R01GM107209. This work used resources of the Northwestern University Structural Biology Facility and the Keck Biophysics Facility, supported by NCI-CCSG-P30-CA060553 awarded to the Robert H. Lurie Comprehensive Cancer Center. Use of the Advanced Photon Source was supported by the US Department of Energy, Office of Science, Office of Basic Energy Sciences, under Contract No. DE-AC02-06CH11357. Use of the Pilatus 3 1M detector was provided by grant 1S10OD018090-01 from NIGMS. Use of the LS-CAT was supported by the Michigan Economic Development Corporation and the Michigan Technology Tri-Corridor (085P1000817). This project was supported by grant 9 P41 GM103622 from the National Institute of General Medical Sciences of the National Institutes of Health. The authors declare no competing financial interests.

## Additional Information

Accession Codes: Coordinates and structure factors have been deposited at the wwPDB (PDB) under the accession code 6D71.

## Notes

https://www.rcsb.org/structure/6D71

