## Supplementary Material for "Structural assembly of the human Miro1/2 GTPases based on the crystal structure of the N-terminal GTPase domain"

**Supplementary Table 1.** *Crystallographic Statistics*

| **Data Collection** | **HsMiro1 nGTPase*GTP** |
|---|---|
| Wavelength | 0.979130 |
| Resolution range (Å) | 61.58 - 1.72 (1.75 - 1.72) |
| Space group | P 21 21 21 |
| Unit Cell Dimensions (Å) | 41.35 78.27 99.78 |
| Unit Cell Angles | 90 90 90 |
| Total reflections | 171752 (8561) |
| Unique reflections | 35285 (3438) |
| Multiplicity | 4.9 (4.8) |
| Completeness (%) | 99.9 (98.3) |
| Mean I/sigma(I) | 8.1 (0.9) |
| Wilson B-factor | 18.03 |
| $R_{MERGE}$ | 0.136 (1.532) |
| $R_{MEAS}$ | 0.152 (1.715) |
| $R_{PIM}$ | 0.067 (0.759) |
| **Refinement** | |
| Reflections used in refinement | 35276 (3438) |
| Reflections used for $R_{FREE}$ | 2000 (195) |
| $R_{WORK}$ | 0.190 (0.336) |
| $R_{FREE}$ | 0.220 (0.316) |
| Number of non-hydrogen atoms | 3277 |
| macromolecules | 2827 |
| ligands | 66 |
| solvent | 384 |
| Protein residues | 341 |
| RMS(bonds) | 0.004 |
| RMS(angles) | 0.800 |
| Ramachandran favored (%) | 98.5 |
| Ramachandran allowed (%) | 1.5 |
| Rotamer outliers (%) | 0.0 |
| Clashscore | 5.34 |
| Average B-factor ($Å^2$) | 23.33 |
| macromolecules | 22.60 |
| ligands | 13.96 |
| solvent | 30.29 |

Values in parentheses correspond to the highest resolution shell.

**Supplementary Table 2**: *SAXS Parameters*

| Dataset | HsMiro1 | HsMiro2 |
|---|---|---|
| **Rg (Å)** | 37.5 ± 0.9 | 34.7 ± 0.5 |
| **Dmax (Å)** | 131 | 124 |
| **Molecular Weight from Porod Volume (kDa)** | 67.8 | 58.6 |
| **Theoretical MW (kDa)** | 68.7 | 66.0 |
| **Total estimate from GNOM** | 0.850 (Good) | 0.864 (Good) |
| **NSD of DAMMIF reconstructions** | 0.62 ± 0.03 | 0.63 ± 0.02 |
| **DAMMIF $\chi^2$** | 0.086 | 0.078 |

Rg and error in Rg were calculated from weighted linear regression analysis of log(I) vs. $q^2$ from raw SAXS data. Reported Dmax values were determined using GNOM (Svergun, 1992). In determining Dmax, GNOM gives a "total estimate", which identifies common artifacts encountered by the GNOM method. Good/reasonable total estimates indicate that commonly observed errors were not made in determining Dmax for our datasets. The molecular weight was estimated by dividing the Porod volume by 1.7 (Petoukhov et al., 2012). The normalized spatial discrepancy (NSD) is a measure of the quantitative similarity between the different independent runs of DAMMIF (Franke and Svergun, 2009). The reported NSD values indicate that the DAMMIF reconstructions are very stable. $\chi^2$ values compare theoretical scattering data based on DAMMIF reconstructions with the raw data. The relatively large errors in the raw data mean that the reconstructions are extremely likely to accurately represent our data; in other words, over half of the theoretical scattering data falls within the error bars in the raw data. Therefore the $\chi^2$ values are very low. These were calculated as described in (Svergun et al., 1995). DAMMIF reconstructions yielded a good model of the SAXS data throughout the entire q range.

**Supplementary Figure 1:** *Comparison of HsMiro nGTPase and cGTPase*

(A) Sequence alignment of the nGTPase and cGTPase switch regions. Completely conserved sequences in each of the motifs are boxed. Note that the alignment of Sw1 between the nGTPase of Miro and Gem1p is problematic (see text). Ser59 of Sw2 is also highlighted. Note the substantial divergence of Sw1 sequence in the cGTPase domain of Miro and Gem1p.

**nGTPase**

```
              G-1          Sw1        Sw2         G-4     G-5

MIRO1_HUMAN   GEPRVGKTS    FPEEVPP    IVDYSEAE    NKSD    CSAK
MIRO2_HUMAN   GEAQVGKTS    FPEEVPP    IVDYSEAE    NKSD    CSAK
MIRO_DROME    GDAGVGKTS    YPEEVPP    IVDFSAVE    NKID    CSAK
GEM1_YEAST    GDEGVGKSS    FIPTIQD    LIDTSDSD    NKCD    TSAK
GEM1_SCHPO    GDQGVGKSS    NVTSIPK    LVDTQSDS    NKSE    CSAL
              *:  ***:*       :       : *  ..     ** :     **
```

**cGTPase**

```
              G-1          Sw1          Sw2         G-4     G-5

MIRO1_HUMAN   GVKNCGKSG    RQKKIREDHK   LHDISESE    AKSD    FTCN
MIRO2_HUMAN   GARGVGKSA    HQDT-REQP-   LCEVGTDG    SKAD    FSCA
MIRO_DROME    GPKGSGKTG    IGKEFKTN--   LRDIDVRH    TKCD    FSLK
GEM1_YEAST    GKPCCGKSS    EEYSPTIKP-   LQELGEQE    SKAD    ISSR
GEM1_SCHPO    GSKSCGKTA    NTNR--LTP-   LSEIGETD    TKAD    ISTA
              *    **:.                 *  ::.      :*.*    ::
```

(B) Structural comparison of HsMiro1 nGTPase and cGTPase domains. The two structures have been globally superposed. The core-GTPase fold is relatively well conserved. However, the surface representation illustrates the difference in the relationship of the position of the Switch 1 region with respect to the bound nucleotide in the two domains (nGTPase (left), and the cGTPase (right)). The bound GTP molecule (nGTPase) and GMPPCP molecule (cGTPase) are represented as spheres. The G-2/Switch 1 peptides (at the bottom of the image) are colored light blue. In the nGTPase both the G-2/Switch 1 and G-3/Switch 2 are more tightly engaged with the bound nucleotide than in the cGTPase domain. The pointer indicates the relative displacement of the Switch 1 backbone atoms towards the nucleotide in the HsMiro1 nGTPase.

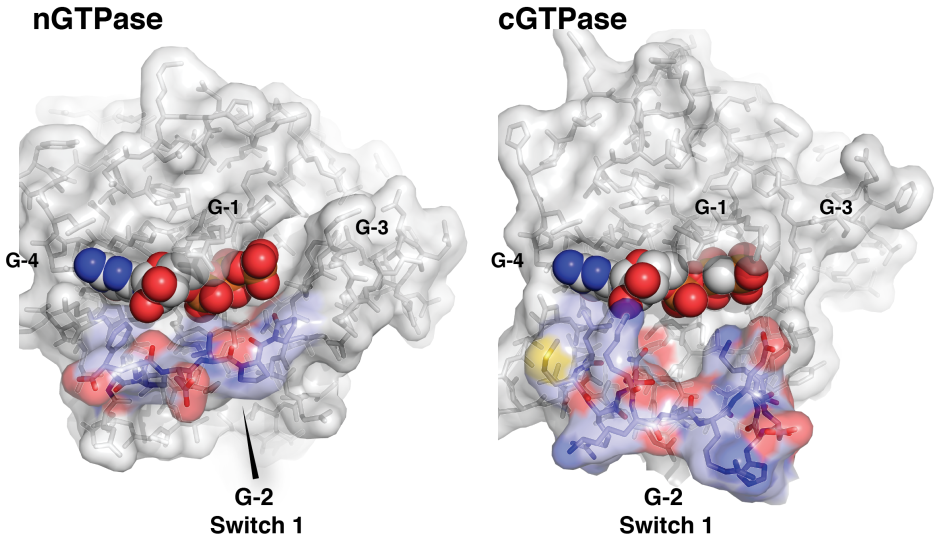
nGTPase
G-1
G-3
G-4
G-2
Switch 1
cGTPase
G-1
G-3
G-4
G-2
Switch 1

**Supplementary Figure 2.** *Interactions between the bound GTP ligand and HsMiro1 nGTPase.*

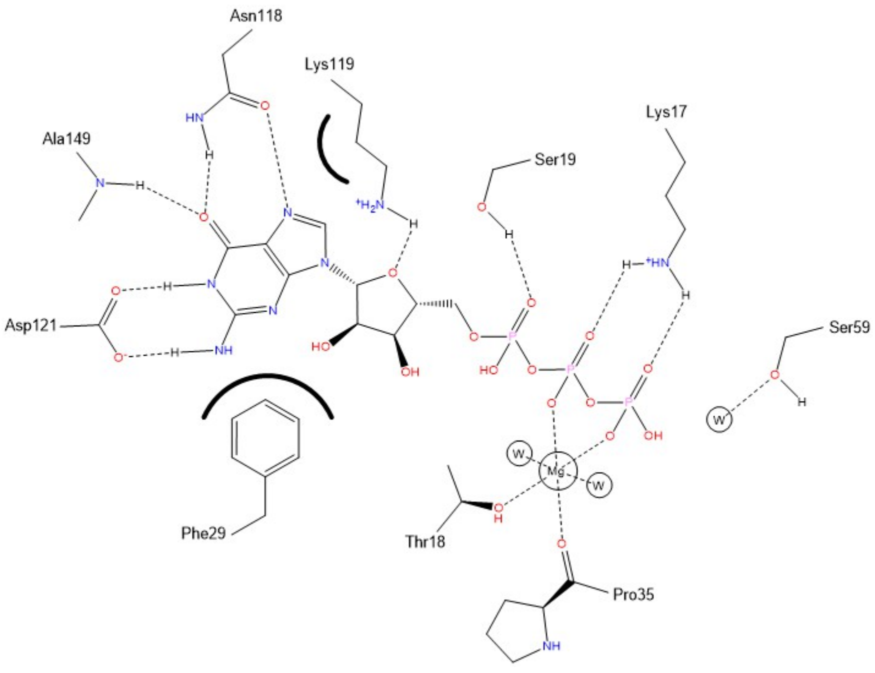

**Supplementary Figure 3:** *SEC-SAXS chromatograms from HsMiro1 and HsMiro2*

The SEC-SAXS elution scattering profiles for HsMiro1 and HsMiro2. The major peak in each sample corresponds to the monomeric Miro species, with average Rgs of 37.5 +/- 0.9 A (HsMiro1) and 34.7 +/- 0.5 A (HsMiro2). Guinier analysis was used for each fraction to obtain both the zero-angle x-ray intensity (blue line, y-axis on left) and the radius of gyration (red line, y-axis on right). Fractions for which the x-ray scattering data was too weak to be extrapolated to an $I_0$ value are not shown. The ranges displayed in this figure were also those used to determine the Dmax and envelopes shown in **Suppl. Fig. 4**. Fraction number is arbitrary. An early eluting peak arises from an uncharacterized and likely artifactual aggregated or disulfide-linked species; multiple runs at different protein concentrations were not consistent with equilibrium binding behavior, and this peak was not analyzed further. Under similar conditions the *Drosophila* DmMiro is monomeric (Klosowiak et al., 2013). A late eluting peak with small size and weak scattering is a buffer species originating during protein preparation.

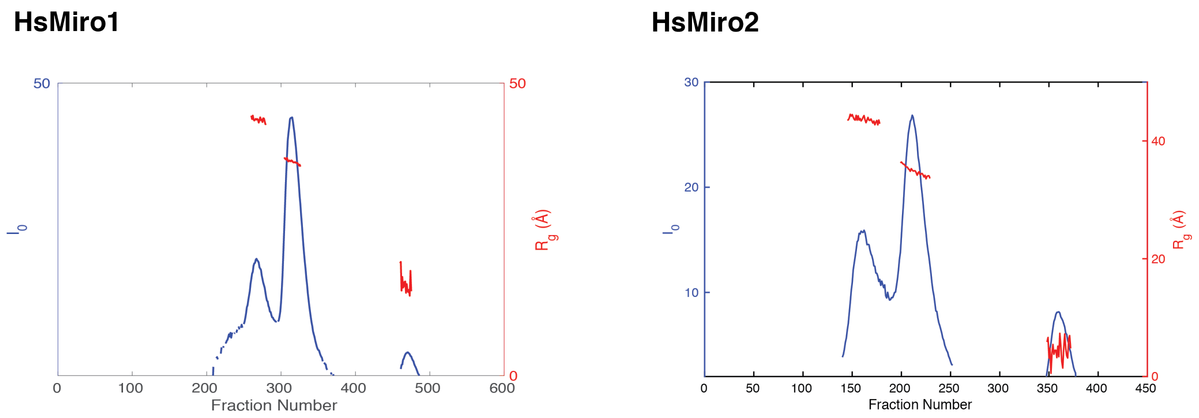

**Supplementary Figure 4:** *SEC-SAXS Analysis of HsMiro1 and HsMiro2.*

**Top,** Log(Intensity) vs. scattering angle. Raw data shown as black dots is compared to the predicted scattering intensity based on the DAMMIF reconstruction in blue. **Second from top**, Guinier plot with linear fit for radius of gyration (*Rg*) measurement. The Guinier analysis was done from the smallest stable angular reading out to a maximum q of1.3 * Rg. **Third from top**, pair distance distribution function *p(r)* indicating maximum dimension (*Dmax*). **Bottom**, Kratky plot showing raw data as dots compared to the regularized intensity obtained by GNOM. This intensity line provided the input data for determining the pair distance distribution function above.

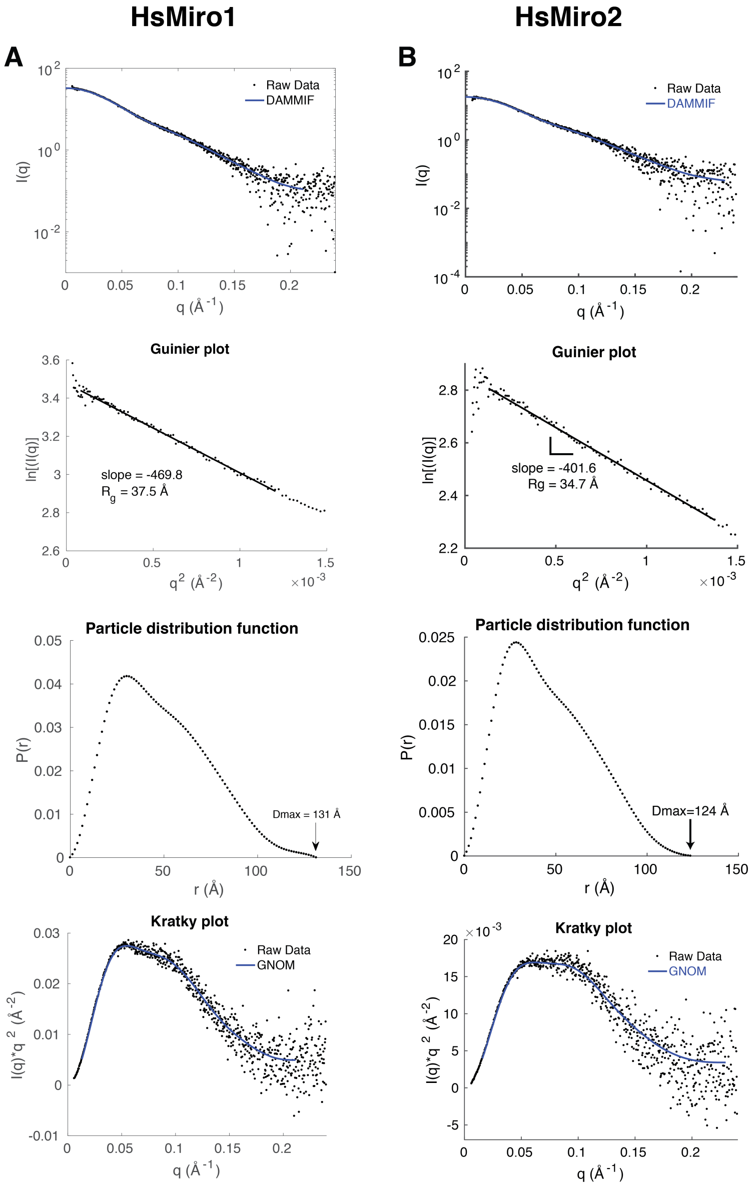
